# Roles of the ClC Chloride Channel CLH-1 in Food-associated Salt Chemotaxis Behavior of *C. elegans*

**DOI:** 10.1101/2020.02.16.951368

**Authors:** Chanhyun Park, Yuki Sakurai, Hirofumi Sato, Shinji Kanda, Yuichi Iino, Hirofumi Kunitomo

## Abstract

The ability of animals to process dynamic sensory information facilitates foraging in an ever changing environment. However, molecular and neural mechanisms underlying such ability remain elusive. The ClC anion channels/transporters play a pivotal role in cellular ion homeostasis across all phlya. Here we find a ClC chloride channel is involved in salt concentration chemotaxis of *C. elegans*. Genetic screening identified two altered-function mutations of *clh-1* that disrupt experience-dependent salt chemotaxis. Using genetically encoded fluorescent sensors, we demonstrate that CLH-1 contributes to regulation of chloride and calcium dynamics of salt-sensing neuron, ASER. The mutant CLH-1 reduced responsiveness of ASER to salt stimuli in terms of both temporal resolution and intensity, which could disrupt navigation strategies for approaching preferred salt concentration. Furthermore, other ClC genes appeared to act redundantly in salt chemotaxis. These findings provide insights into the regulatory mechanism of neuronal responsivity by ClCs that contribute to modulation of navigation behavior.

## Introduction

Memorizing environmental conditions associated with food and generating an optimal foraging strategy based on those memories are basic and important abilities for survival. Mechanisms of food-associated learning have long been addressed in many species, dating back to Pavlovian appetitive conditioning demonstrated in dogs (Braubach et al., 2009; Cho et al., 2016; Gottfried et al., 2003; Hirano et al., 2013; O’Doherty et al., 2003; Otis et al., 2017; Pavlov, 1927; Sasakura and Mori, 2013; Winter and Stich, 2005). By virtue of its simple nervous system and amenability to genetic manipulations, the soil nematode *Caenorhabditis elegans* has been used to unveil molecular and neural mechanisms of learning. *C. elegans* shows food-associated learning in combination with various sensory modalities including gustatory, olfactory, thermosensory, and mechanosensory cues (Colbert and Bargmann, 1995; Hedgecock and Russell, 1975; Kindt et al., 2007; Saeki et al., 2001). We have previously reported that *C. elegans* shows plasticity in chemotaxis to salt (sodium chloride; NaCl); wild-type animals are attracted to the salt concentration at which they have been fed, while avoiding concentrations at which they have been starved (salt concentration chemotaxis, Kunitomo et al., 2013, Figure 1**a**). *C. elegans* senses inorganic ions mainly through the bilateral salt-sensing neuron pair, ASE (Bargmann and Horvitz, 1991). Sensory input to ASE-right (ASER), is essential and sufficient for food-associated salt chemotaxis. Modulation of synaptic transmission from ASER to the first layer interneurons AIA, AIB and AIY, whose activity regulates exploratory behaviors (Gray et al., 2005; Li et al., 2014; Piggott et al., 2011), is also implicated in modification of salt chemotaxis (Kunitomo et al., 2013; Luo et al., 2014; Wang et al., 2017).

**Figure 1.**
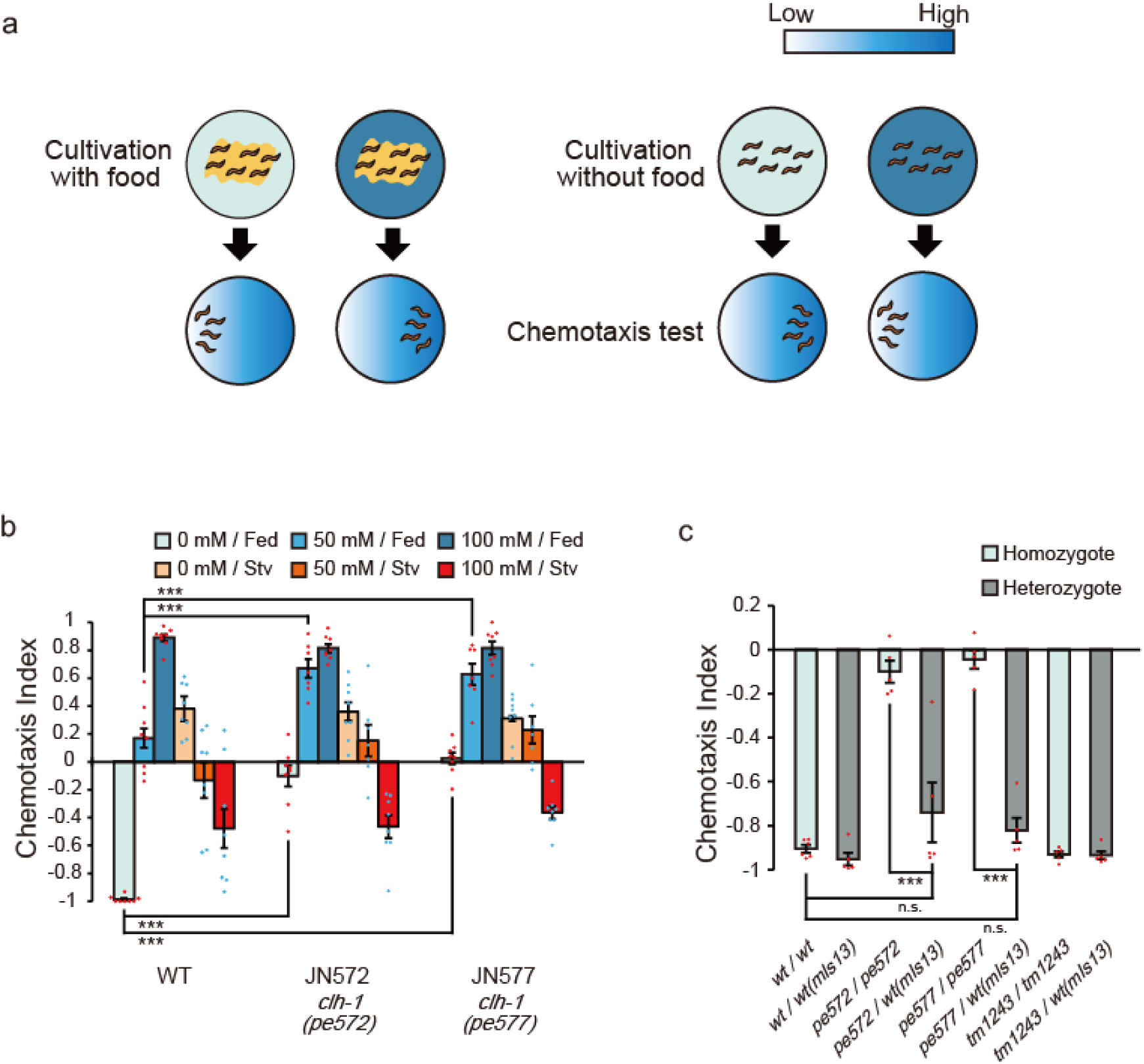
Two missense mutations in *clh-1* give rise to food-associated salt chemotaxis disorder. (**a**) Salt concentration chemotaxis. Animals were cultivated at 0 mM or 100 mM of NaCl with or without food and placed on an assay plate with NaCl gradient. Distribution of animals was quantified by calculating a chemotaxis index. See method for details. (**b**) Chemotaxis of wild-type animals and two mutants obtained from screening, JN572: *clh-1(pe572)* and JN577: *clh-1(pe577)*. Dots show the result of individual trials. Bar and the error bar represent mean +/− s.e.m., *n* = 8 assays, Dunnett’s test, \*\*\**p* < 0.001, n.s., not significant. (**c**) Chemotaxis of *clh-1* heterozygotes. *clh-1* homozygotes were crossed with a *clh-1(wt)* reporter strain that express GFP in pharyngeal muscle (*mIs13*). Resulted F1 animals were used for assay. Dots show the result of individual trials. Bar and the error bar represent mean +/− s.e.m., *n* ≧ 4, Tukey’s test, \*\*\**p* < 0.001.

Molecular mechanisms that regulate perception and propagation of salt stimuli in ASER have been proposed. A receptor-type guanylyl cyclase GCY-22 plays a pivotal role in perception of salt stimuli and is suggested to act as an ion receptor of ASER (Adachi et al., 2010; Ortiz et al., 2009; Smith et al., 2013). Excitation of ASER depends also on cyclic nucleotide-gated (CNG) channels consisting of TAX-2 and TAX-4 (Suzuki et al., 2008). *C. elegans* genome does not contain typical voltage-gated sodium channel genes (Bargmann, 1998). Instead, voltage-gated calcium channels are responsible for propagation of depolarization, which is examined by electrophysiological studies (Goodman et al., 1998; Shindou et al., 2019). However, contribution of anions to the regulation of ASER activity has not been discussed yet.

Anion transporters play critical roles in regulating excitability of neurons as they finely tune electrophysiological properties of membranes. However, how and which molecules contribute to the responsivity of specific neurons and eventually produces behavioral output, remains rudimentary. Here, we identify the ClC chloride channel CLH-1 as a possible regulator of food-associated salt chemotaxis in *C. elegans*. ClC proteins transport univalent anions across membranes to control electrochemical potential of excitable cells and to maintain ionic milieu as well as pH of intracellular organelles (Ahnert-Hilger and Jahn, 2011; Bösl et al., 2001; Branicky et al., 2014). Malfunction of ClC genes results in various diseases such as myotonia, leukodystrophy, hyperaldosteronism and epilepsy in humans (Blanz et al., 2007; Charlet-B. et al., 2002; Fernandes-Rosa et al., 2018; Yamamoto et al., 2015). The causal relationship among ClC malfunction, physiological consequences, and disease manifestations are not fully understood in many cases. CLH-1 shares the highest (37%) identity with mammalian ClC-2. Functional studies using heterologous expression in *Xenopus* oocytes and mammalian cells demonstrated that both CLH-1 and ClC-2 are inwardly rectifying chloride channels (Grant et al., 2015; Nehrke et al., 2000; Staley et al., 1996; Thiemann et al., 1992). On the other hand, two recent studies showed that ClC-2 contributes to Cl^−^ influx in neurons depending on the electrochemical potential of Cl^−^ (Ratté and Prescott, 2011; Rinke et al., 2010). Thus, the function of CLH-1/ClC-2 in the nervous system remains elusive.

In this study, we show two novel missense mutations in *clh-1* change NaCl concentration preference of *C. elegans* only after food experience. Genetic analyses revealed that CLH-1 acts in ASER and that both appropriate quantitative and spatial function of CLH-1 is required for normal low-salt chemotaxis. Functional imaging of neurons indicated that mutations in *clh-1* altered the responsivity of ASER and AIB neurons, which may consequently affect behavioral outputs. These results suggest that responsivity of the salt circuit is maintained by CLH-1 to generate proper navigation behavior in salt chemotaxis.

## Results

### Missense mutations of *clh-1* give rise to a disorder in food-associated salt concentration chemotaxis

*C. elegans* is attracted to the salt concentration at which they have been fed, while they avoid concentrations at which they have been starved (Figure 1**a** and Figure 1—figure supplement 1**a**). To better understand the molecular mechanisms of salt chemotaxis plasticity, we screened for mutants that showed defects in salt chemotaxis after feeding but not after starvation (See Materials and methods, Figure 1—figure supplement 1**b**). Two mutants, JN572 and JN577, showed a similar phenotype: impaired chemotaxis toward low salt after feeding on NaCl-free nematode growth medium (NaCl-free NGM, hereinafter referred to as cultivation at 0 mM NaCl). Chemotaxis to high salt after feeding on NGM with a salt concentration of 100 mM NaCl (hereinafter referred to as cultivation at 100 mM NaCl), and chemotaxis after starvation were comparable to those of wild-type animals (Figure 1**b** and Figure 1—figure supplement 1**c**). Neither shortened nor extended cultivation at 0 mM NaCl did not ameliorate the defects (Figure 1—figure supplement 1**d**, **e**), suggesting that the impaired chemotaxis is not due to the delay of behavioral modification. Rather, the mutants are unable to generate migration bias toward low salt. Consistent with this idea, the mutants showed a preference for high salt after cultivation at 50 mM NaCl, under which condition wild-type animals showed unbiased salt preference for either low or high salt concentration (Figure 1**b**). Taken together, JN572 and JN577 show a defect in chemotaxis toward low salt after cultivation in the presence of food.

We mapped mutations of JN572 and JN577 between genetic positions 2.82 and 6.12 (cM) on chromosome II (See Materials and methods, Fay and Bender, 2006; Wicks et al., 2001). Genome sequencing revealed that they carried distinct missense mutations in the *clh-1* gene, one of six ClC channels/transporters in *C. elegans*. Mutations predicted M293I and I146T substitutions in CLH-1A in JN572 and JN577, respectively, and they are hereinafter referred to as *clh-1(pe572)* and *clh-1(pe577)* (Figure 1—figure supplement 2**a**, **b**). Salt chemotaxis defects of the mutants were rescued by a *clh-1(wt)* genomic fragment, confirming that *clh-1* is the responsible gene (Figure 1—figure supplement 2**c**). We also noticed that extra copies of *clh-1(wt)* genomic fragment weakened low-salt chemotaxis of wild type, implying that overexpression of CLH-1 could disrupt chemotaxis to low salt (Figure 1—figure supplement 2**c**).

Interestingly, deletion mutants of *clh-1*, all of which harbor a lesion in the pore-forming transmembrane domain of CLH-1 and hence are putative loss-of-function alleles, showed almost no discernible defect in salt chemotaxis (Figure 1—figure supplement 2**a**, 3**a**). This result suggests that chemotaxis defects of *clh-1(pe572)* and *clh-1(pe577)* were caused by anomalous activity of *clh-1*. To characterize the nature of *clh-1* missense alleles, we observed food-associated salt chemotaxis of heterozygotes. *clh-1(pe572)/wt* and *clh-1(pe577)/wt* showed normal low-salt chemotaxis, demonstrating that both missense alleles are recessive to wild-type allele (Figure 1**c**). Also, we noted that the effects of both missense alleles on salt chemotaxis are dosage-dependent, that is, *clh-1(pe572)/clh-1(tm1243)* and *clh-1(pe577)/clh-1(tm1243)* showed a modest defect in salt chemotaxis to low salt after cultivation at 0 mM NaCl in contrast to the strong defect of *clh-1(pe572)/clh-1(pe572)* and *clh-1(pe577)/clh-1(pe577)* (Figure 1—figure supplement 3**b**). Furthermore, we unexpectedly found that *clh-1(tm1243)* conferred a weak resistance to an acetylcholine receptor agonist, levamisole. On the contrary, *clh-1(pe572)* or *clh-1(pe577)* caused an enhanced sensitivity to levamisole if compared with wild type, suggesting these alleles may not be simple reduction-of-function alleles (Figure 1—figure supplement 3**c**). Together, these data show that *clh-1(pe572)* and *clh-1(pe577)* (hereinafter collectively referred to as *clh-1(pe)*) are recessive neomorphic mutations whose salt chemotaxis phenotype appears in a dosage-dependent manner.

Given that salt chemotaxis may be affected by the dosage of *clh-1(pe)* alleles, we wondered if overexpression of *clh-1(pe)* can give rise to salt chemotaxis defect. To examine this possibility, we introduced either *clh-1(pe)* genomic DNA fragments into wild type or *clh-1(tm1243)* mutants. Overexpression of mutant *clh-1* conferred defects in low-salt chemotaxis in both genomic backgrounds, demonstrating that excess *clh-1(pe)* override the canonical *clh-1(wt)* function (Figure 1—figure supplement 3**d**). Therefore, excess *clh-1*, either wild-type or *pe* alleles, can impair low-salt chemotaxis. In conclusion, *clh-1(pe572)* and *clh-1(pe577)* are neomorphic alleles that disrupt chemotaxis to low salt after cultivation at 0 mM NaCl.

### ClC proteins redundantly act in salt chemotaxis

All ClC anion channels/transporters characterized so far function as either homodimers or heterodimers (Accardi, 2015; Stölting et al., 2014a). This raised the possibility that mutant CLH-1 molecules impaired the function of other CLH gene products by forming heterodimers with them, thereby resulting in defective salt chemotaxis. To examine this, we observed salt chemotaxis of the mutants whose *clh* genes were deleted individually or in combinations. The *C. elegans* genome carries 6 ClC genes (*clh-1* through *clh-4* for channel, and *clh-5* and *clh-6* for transporter, predicted from a key amino acid residue and subcellular localization, Figure 2—figure supplement 1**a**, Schriever et al., 1999; Nehrke et al., 2000). Each single mutant except for *clh-5(tm6008)* showed normal salt chemotaxis under fed conditions (Figure 2**a**, Figure 2—figure supplement 1**b**). For multiple mutants, we started from *clh-2(ok636) clh-1(tm1243)* double mutants because they had the highest homology (52% identity) among *clh* genes. Although the double mutant showed no chemotaxis defect, triple mutants with either *clh-3(ok763)* or *clh-6(tm617)* showed impaired chemotaxis toward high salt regions, indicating that they act redundantly with *clh-1* and *clh-2* (Figure 2**b**, Figure 2—figure supplement 1**c**). Triple mutants *clh-3(ok763) clh-2(ok636) clh-1(tm1243)* and *clh-5(tm6008) clh-2(ok636) clh-1(tm1243)* showed a weak defect in chemotaxis to low salt. However, the phenotype of *clh-1(pe572)* and *clh-1(pe577)* mutants was not recapitulated in any tested mutants. Interestingly, the defect of *clh-5(tm6008) clh-2(ok636) clh-1(tm1243)* were partially restored in *clh-3(ok763) clh-5(tm6008) clh-2(ok636) clh-1(tm1243)* quadruple mutants (Figure 2**b**). These results suggest that the chemotaxis defect of *clh-1(pe)* mutants is unlikely to be due to impairment of other CLH proteins.

**Figure 2.**
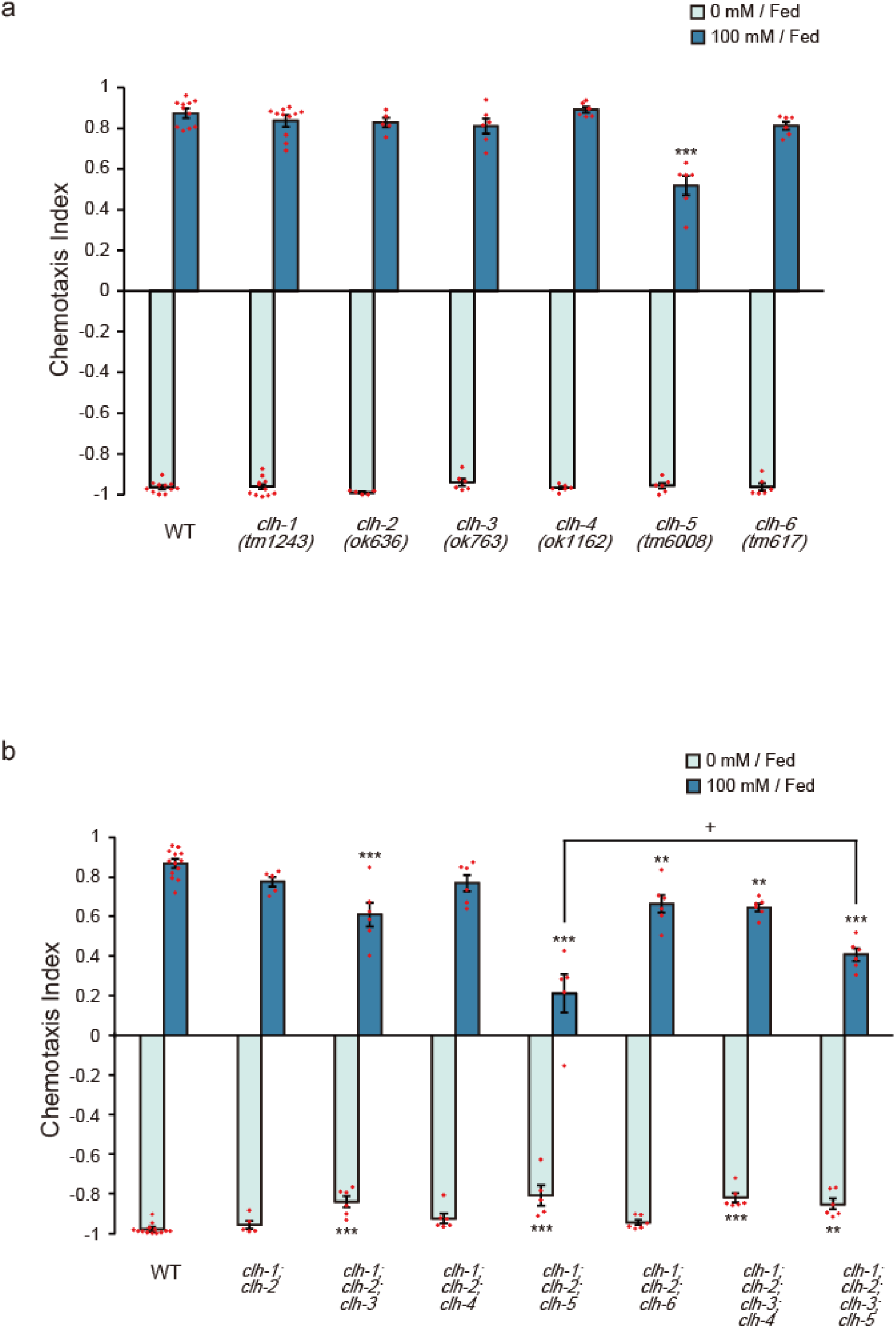
ClC genes function redundantly in salt chemotaxis. (**a**) Chemotaxis of deletion mutants of 6 ClC genes (*clh-1* to *clh-6*). All mutant strains were outcrossed with wild type more than 4 times. Dot shows the result of individual trials. Bar and the error bar represent mean +/− s.e.m., *n* ≧ 5 assays, compared with wild type, Dunnett’s test, \*\*\**p* < 0.001. (**b**) Chemotaxis of *clh* multiple mutants. Dot shows the result of individual trials. Bar and the error bar represent mean +/− s.e.m., *n* ≧ 5 assays, Tukey’s test, \*\*\**p* < 0.001, \*\**p* < 0.01, \**p* < 0.05, compared with wild type. +*p* < 0.05, compared between indicated mutants.

### CLH-1 acts in ASER to affect salt preference after cultivation at 0 mM NaCl

It has previously been reported that *clh-1* is expressed in hypodermal cells, seam cells, D-cells of the vulva, and neuronal and glial cells of the head (Grant et al., 2015; Nehrke et al., 2000). These expression patterns were confirmed with a transcriptional reporter (*clh-1p::nls4::mTFP*, see methods). Of the head neurons, we found that at least ASE, AWA, and AWC sensory neurons expressed the reporter (Figure 3**a**). To determine the site of action of *clh-1*, we performed cell-specific rescue experiments using *clh-1(wt)* cDNA. The mutant phenotype was rescued when cDNA was expressed either pan-neuronally or specifically in ASER, suggesting that *clh-1* acts in the nervous system including ASER (Figure 3**b**, similar results for *pe577*, Figure 3—figure supplement 1**a**). On the other hand, *clh-1(wt)* cDNA failed to rescue *clh-1(pe572)* when expressed either in amphid sheath (AmSh) cells, where CLH-1 function as a pH mediator (Grant et al., 2015), or in the left-sided ASE neuron (ASEL) (Figure 3**b**). Unexpectedly, combined expression of the transgene in ASER and AmSh cells or expression in all ciliated neurons also failed to rescue the phenotype (Figure 3**b**), albeit it recovered chemotaxis of very limited population of animals. These phenotypes are reminiscent of the weakened low-salt chemotaxis by overexpression of *clh-1(wt)* genomic fragment (Figure 1—figure supplement 2**c**), indicating a possibility that excessive function of CLH-1*(wt)* in the tissue near ASER could disturb low-salt chemotaxis.

**Figure 3.**
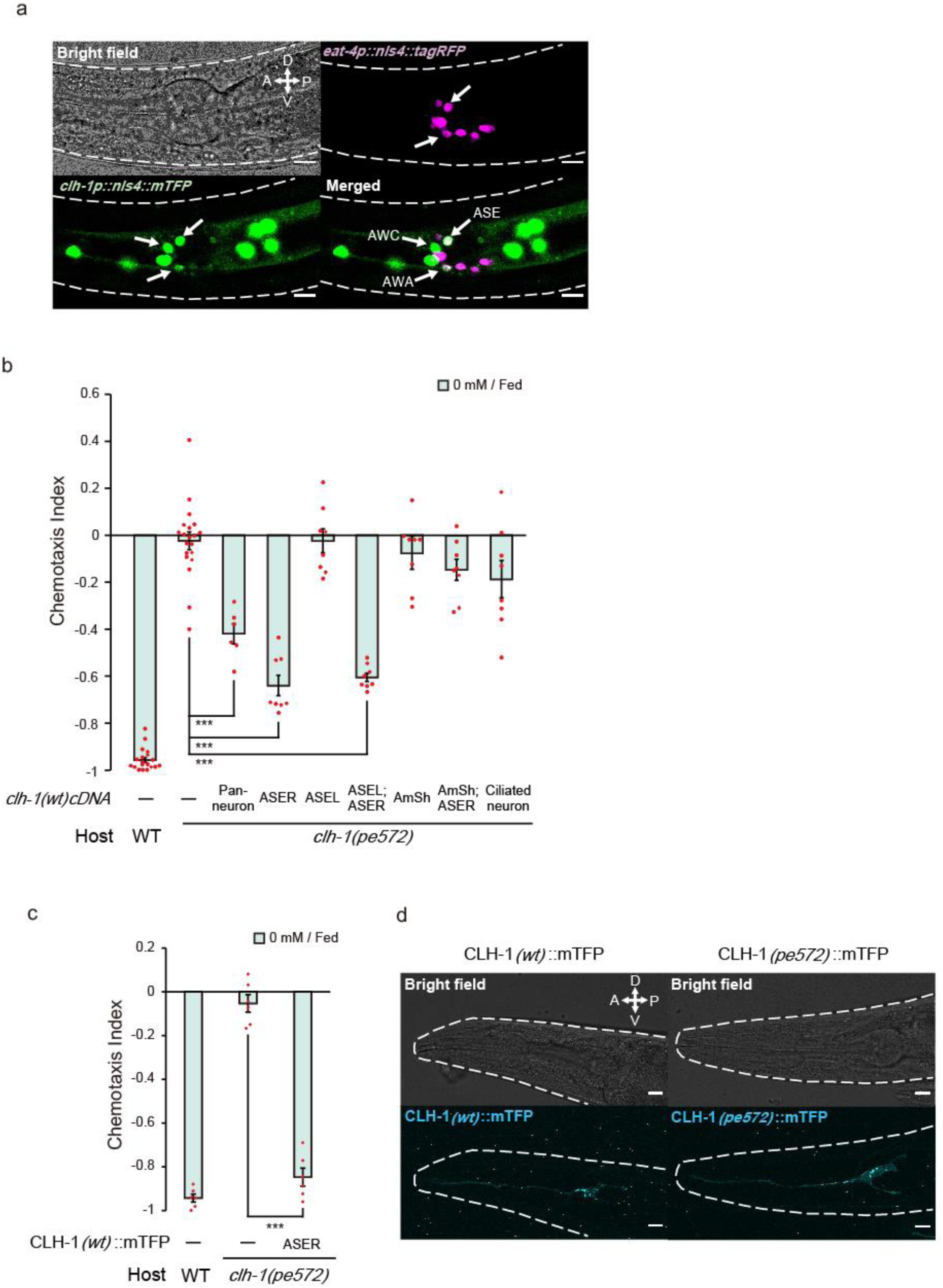
*clh-1* acts in the salt sensing neuron ASER. (**a**) Expression pattern of *clh-1p::nls4::mTFP* (green) in an adult animal that express glutamatergic neuron-specific marker *eat-4p::nls4::tagRFP* (magenta). Three pairs of sensory neurons, namely, AWA, AWC and ASE expressed *clh-1p::nls4::mTFP*. DiI, which stain 6 pairs of head sensory neurons, was also used as a position marker for cell identification (data not shown). Scale bar = 10 µm. (**b**) Rescue of *clh-1(pe572)* mutants by cell-specific expression of *clh-1(wt)* cDNA. Promoters used in this experiment are as follows; *rimb-1p* for all neurons, *gcy-5p* for ASER, *gcy-7p* for ASEL, *vap-1p* for amphid sheath cells, *dyf-11p* for ciliated neurons. Dot shows the result of individual trials. Bar and the error bar represent mean +/− s.e.m., *n* ≧ 6 assays, Tukey’s test. \*\*\**p* < 0.001. (**c**) Chemotaxis of *clh-1(pe572)* mutants that express *clh-1(wt)cDNA::mTFP* in ASER. mTFP-tagged CLH-1 is functional. Dot shows the result of individual trials. Bar and the error bar represent mean +/− s.e.m., *n* = 6 assays, Tukey’s test. \*\*\**p* < 0.001. (**d**) Expression of *gcy-5p::clh-1(wt)cDNA::mTFP* in wild type (left) and *gcy-5p::clh-1(pe572)cDNA::mTFP* in *clh-1(pe572)*. Both CLH-1*(wt)*::mTFP and CLH-1*(pe572)*::mTFP fluorescence were observed in membrane of dendrite, soma, axon and cell organelles. Scale bar = 10 µm.

### Morphology of ASER and localization of CLH-1 are largely unaffected by missense mutations

Head sensory neurons of *C. elegans* sense environmental stimuli via neural receptive endings, which are comprised of cilia and microvilli (Perkins et al., 1986; Ward et al., 1975; Ware et al., 1975). Function of these endings largely depends on glia which ensheathe them. A recent study showed that AmSh glia regulate the function and shape of AFD thermosensory neuron’s microvilli by modulating efflux of Cl^-^ to the extracellular space of the receptive endings (Singhvi et al., 2016). In addition, it has been elucidated that CLH-1 transports anions to maintain intracellular pH of AmSh cells (Grant et al., 2015). These results implied the possibility that the receptive ending of ASER may be impaired in *clh-1(pe)* mutants. However, we did not find any notable change in the morphology of ASER including its sensory cilium (Figure 3—figure supplement 1**b**-**d**). Exposure to different salt and food conditions did not affect the length of ASER sensory cilium both in wild type and in *clh-1* mutants, suggesting that the morphology of ASER receptive ending remained largely unchanged (Figure 3—figure supplement 1**e**).

To examine subcellular localization of CLH-1 in ASER, we generated an mTFP-tagged CLH-1 (CLH-1::mTFP). The fusion protein was functional (Figure 3**c**). Fluorescent signals were detected in the plasma membrane and cell organelles (Figure 3**d**, left), as previously reported in mammalian ClC channels and ClC transporters, respectively (Jentsch, 2008, 2007). Localization patterns of CLH-1::mTFP with M293I mutation were comparable to those of wild type (Figure 3**d**, right). These results indicate that the mutation does not affect intracellular localization of CLH-1.

### Salt stimulus changes intracellular chloride concentration of ASER and this response is altered in *clh-1(pe)* mutants

Previous studies showed that CLH-1 mediates flux of univalent anions including chloride and bicarbonate. Furthermore, electrophysiological characterization of CLH-1 using *Xenopus* oocytes indicated that CLH-1 is an inwardly rectifying channel that conduct Cl^-^ (Grant et al., 2015; Nehrke et al., 2000). Our genetic analyses suggested the *clh-1(pe)* mutations might be neomorphic. To examine whether the mutations affected the property of CLH-1 conductance, we expressed mutant CLH-1 in *Xenopus* oocytes and measured membrane currents via two-electrode voltage clamping. Wild-type CLH-1 showed inwardly rectifying currents from −160 to −80 mV as previously reported (Figure 4**a-e**, Grant et al., 2015; Nehrke et al., 2000). Similar currents were also observed in both CLH-1(M293I) and CLH-1(I146T). This result supports the idea that the basic function of CLH-1 as a voltage-dependent anion channel is retained in CLH-1(M293I) and CLH-1(I146T) mutants.

**Figure 4.**
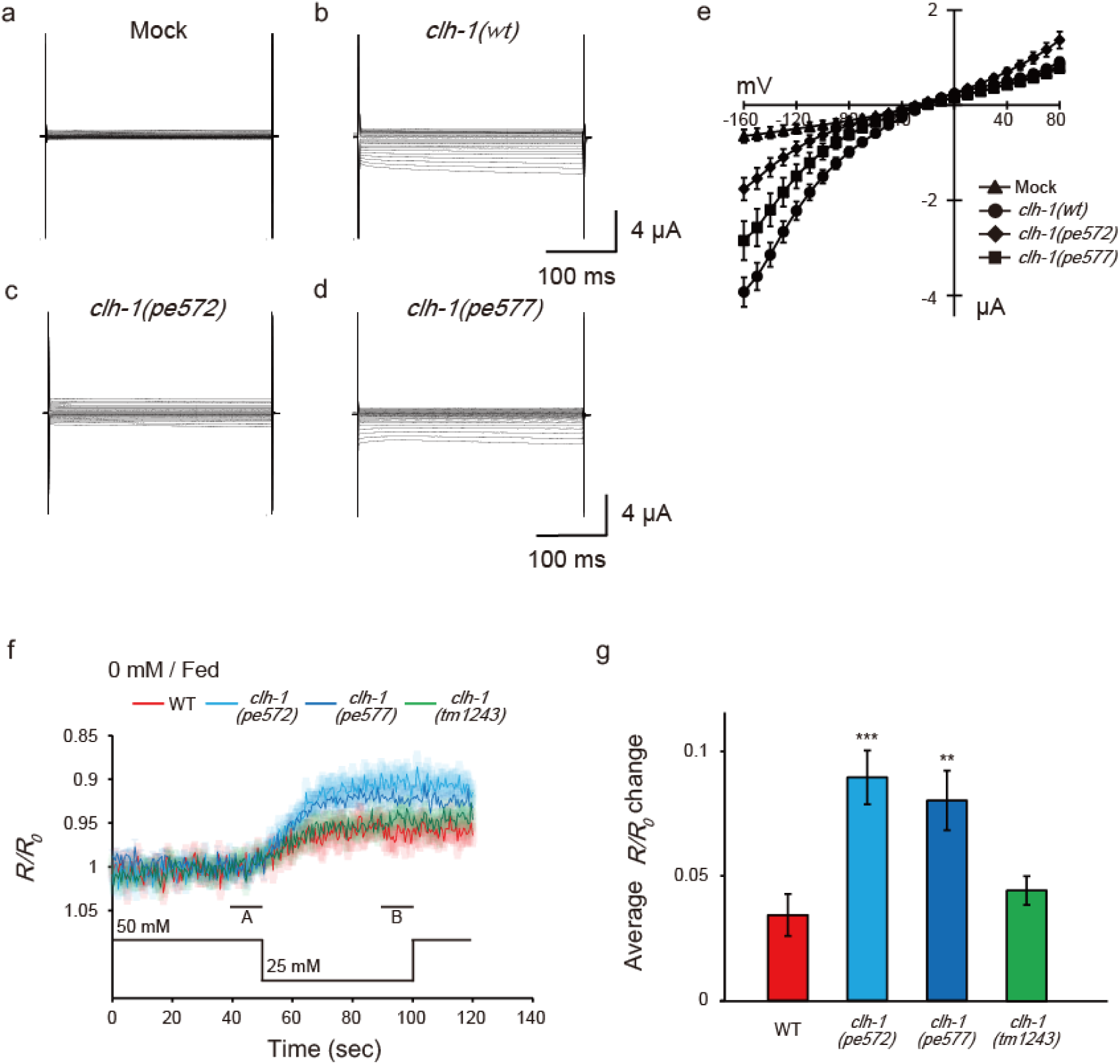
Mutations in *clh-1* change chloride dynamics of ASER in response to salt down-step stimulus. (**a-e**) Representative current traces from *Xenopus* oocytes that express cRNA for mock (**a**), *clh-1(wt)* (**b**), *clh-1(pe572)* (**c**), *clh-1(pe577)* (**d**), clamped at voltages ranging from −160 mV to 80 mV. (**e**) The averaged current-voltage relationships of mock (triangle, *n* = 7), *clh-1(wt)* (circle, *n* = 20), *clh-1*(*pe572)* (diamond, *n* = 13) and *clh-1(pe577)* (square, *n* = 16). The error bars represent s.e.m. (**f**) Averaged Superclomeleon responses of ASER after cultivation at 0 mM NaCl in the presence of food. External NaCl concentration was shifted from 50 mM to 25 mM at time 50 s. Note that the scale of the vertical axis is inverted so that increase in chloride concentration is displayed as up-shift of traces. The shaded region represents s.e.m., *n* ≧ 17 animals. (**g**) *R/R_0_* changes upon salt decrease. A and B indicate the time points for calculating *R/R_0_* changes. Mean +/− s.e.m., *n* ≧ 17 animals.

Next, we asked how *clh-1* mutations disturb salt chemotaxis of *C. elegans*. We focused on the salt-sensing neuron ASER, one of the site-of-action of *clh-1*, and which is essential for food-associated salt concentration chemotaxis (Kunitomo et al., 2013). ASER is activated by salt concentration decreases and deactivated by salt concentration increases (Suzuki et al., 2008, see below). Given that CLH-1 localized to the membranous compartments of ASER (Figure 3**d**) and acted as a chloride channel (Figure 4**a-e**), we hypothesized that CLH-1 might be involved in chloride dynamics during salt response of ASER. To examine this possibility, we monitored intracellular chloride ([Cl^-^]_i_) dynamics of ASER by utilizing the genetically encoded chloride indicator, Superclomeleon. This probe is a FRET-type indicator in which the YFP/CFP fluorescence ratio decreases upon binding to chloride ions (Grimley et al., 2013). Animals were cultivated at 0 mM NaCl with food and immobilized in the olfactory chip (Chronis et al., 2007), and a NaCl down-step from 50 mM to 25 mM was applied as salt stimulus. In wild type, YFP/CFP ratio decreased upon salt down-step, indicating that [Cl^-^]_i_ of ASER was increased when the neuron was activated (Figure 4**f**). This response is not, at least solely, mediated by CLH-1 because it was also observed in *clh-1(tm1243)* mutants. Notably, the magnitude of YFP/CFP ratio change was significantly larger in *clh-1(pe)* mutants, suggesting that ASER [Cl^-^]_i_ greatly increased in mutants (Figure 4**g**).

### ASER salt response is altered in *clh-1* mutants

Intracellular chloride concentration increase can antagonize depolarization of neurons. We then wanted to look into whether *clh-1* mutations affect the activity of ASER in response to salt stimulus. As aforementioned, ASER is activated by salt concentration decreases, which is indicated by an increase in intracellular calcium levels, whereas it is deactivated by salt concentration increases (Suzuki et al., 2008). Such ASER responsivity is basically retained regardless of cultivation salt concentrations or food availability (Kunitomo et al., 2013; Oda et al., 2011).

We performed *in vivo* calcium imaging in wild type and *clh-1* mutants using a genetically encoded calcium indicator YC2.6 (Chronis et al., 2007; Horikawa et al., 2010). Animals were cultivated at either 0 mM NaCl or 100 mM NaCl with or without food, and ASER was stimulated by repeated salt concentration changes from 50 mM NaCl to 25 mM NaCl, to observe responses to both down-step and up-step stimuli. After cultivation at 0 mM NaCl, the amplitude of calcium response to the first down-step stimulus was comparable between wild type and *clh-1* mutants. However, the response to the second down-step was diminished in *clh-1(pe)* mutants compared to wild type (Figure 5**a**, **b**). A similar trend was observed in the third down-step response. The decay of intracellular calcium level ([Ca^2+^]_i_) was small in the *clh-1(pe)* mutants, which was more evident after salt up-step (Figure 5—figure supplement 1**a**, **b**). This diminished decay was likely responsible for the decreased calcium response to the repeated stimuli. On the other hand, ASER salt response of *clh-1(tm1243)* was similar to that of wild type, except that the decay was significantly large during the first down-step stimulus (Figure 5**a**, **b** and Figure 5—figure supplement 1**a**, **b**). Interestingly, reduction of ASER response amplitude upon repeated salt down-step stimuli was not obvious after cultivation at 100 mM NaCl with food, although *clh-1(pe572)* constantly showed small ASER responses (Figure 5**c**, **d** and Figure 5—figure supplement 1**c**, **d**). Considering the essential role of ASER in salt concentration chemotaxis, these results imply that hampered chemotaxis of *clh-1(pe)* mutants toward low salt after low-salt cultivation is probably due to the abnormal ASER responsivity to salt concentration change.

**Figure 5.**
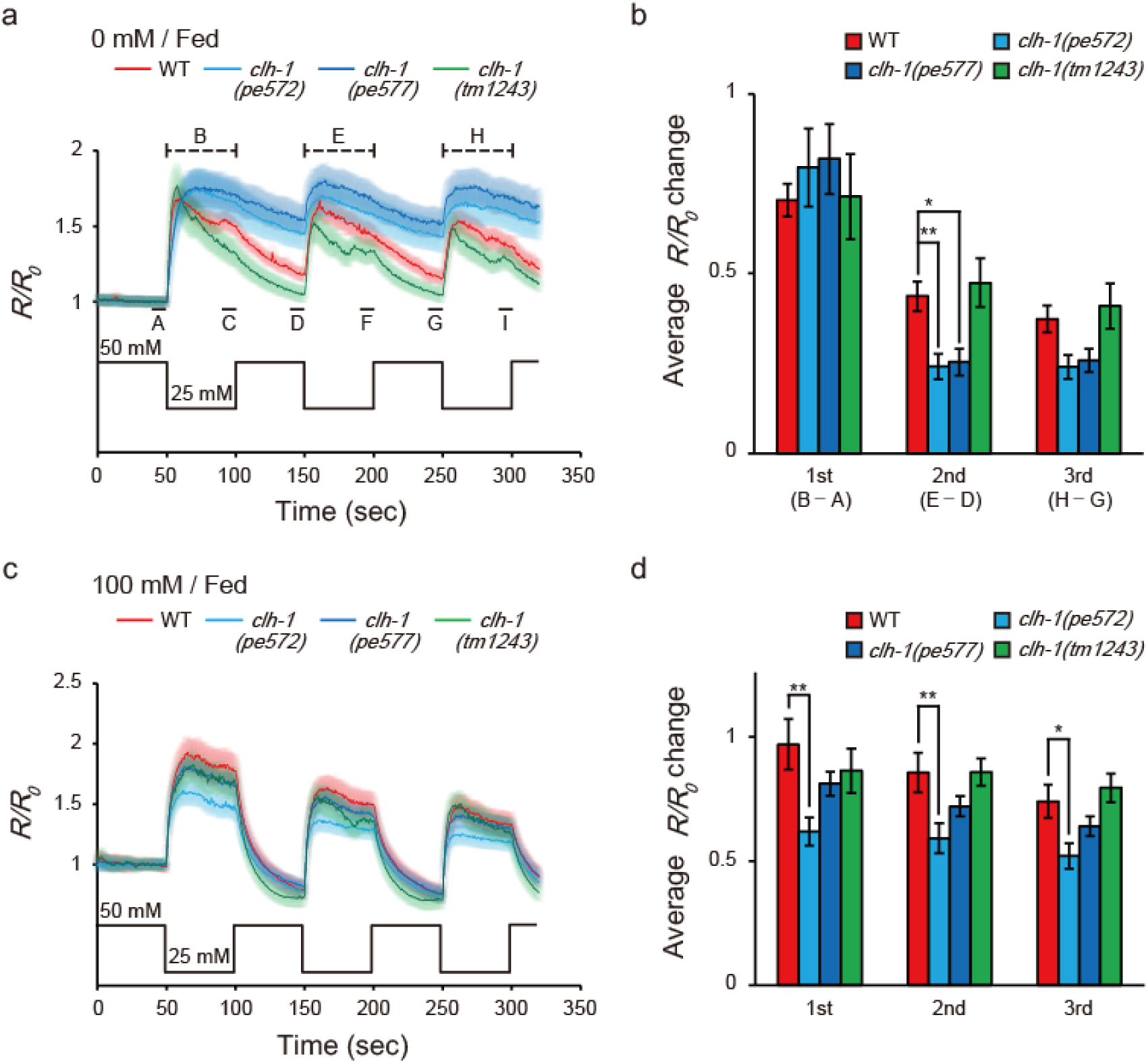
Mutations in *clh-1* change calcium dynamics of ASER in response to repeated salt stimuli. (**a** and **c**) Averaged calcium responses of ASER after cultivation at 0 mM NaCl (**a**) or 100 mM NaCl (**c**) in the presence of food, and stimulated by NaCl concentration changes between 50 mM and 25 mM. A to I indicate the time points for calculation of *R/R_0_* intensity changes. A, D, and G are the time points for pre-stimulus *R/R_0_*, B, E, and H are the time points for peak *R/R_0_* during stimulation, C, F, and I are the time points for decayed *R/R_0_* during stimulation. See methods for details. The shaded region represents s.e.m., *n* ≧ 16 animals. (**b** and **d**) *R/R_0_* changes at each NaCl down-step stimulus (B - A, E - D, and H - G for 1st, 2nd, and 3rd stimulus, respectively). See methods for details. 0 mM NaCl cultivated (**b**) or 100 mM cultivated (**d**). Bar and the error bar represent mean +/− s.e.m., *n* ≧ 16 animals, Dunnett’s test, \*\**p* < 0.01, \**p* < 0.05.

Consistent with previous reports, salt responses of ASER of starved wild-type animals were not largely different when compared to that of fed animals. However, the ASER activity patterns of *clh-1* mutants starved at 0 mM were distinct from those of fed animals (please compare Figure 5, Figure 5—figure supplement 1 and Figure 5—figure supplement 2). After cultivation at 0 mM without food, for example, the amplitude of activation of *clh-1(pe577)* mutants were even larger than that of wild type (Figure 5—figure supplement 2**a**-**c**). Meanwhile, the difference of ASER salt responses between wild type and *clh-1(pe)* mutants were less obvious after starvation at 100 mM NaCl (Figure 5—figure supplement 2**e-h**). Taking all these into account, we concluded that mutations in *clh-1* affect responsivity of ASER, most notably in the reduced response to repeated salt down-step and up-step stimuli after cultivation at 0 mM NaCl with food.

### Behavioral strategies for chemotaxis are disrupted in *clh-1(pe)* mutants

Next, we quantitatively analyzed the navigation behavior of *clh-1* mutants to examine which behavioral components are affected. *C. elegans* utilize at least two behavioral strategies to achieve salt chemotaxis: klinokinesis and klinotaxis. In klinokinesis, migration bias is generated by controlling the frequency of steep turns called pirouettes, which are typically accompanied by reversals and omega turns. The bout of pirouette is triggered according to cumulative salt concentration change along an animal’s progression (Pierce-Shimomura et al., 2001). In klinotaxis, animals gradually curve towards their preferred direction by sensing fluctuation of salt concentration accompanying head bending (Iino and Yoshida, 2009). Input to ASER is both required and sufficient for fed animals to generate the two behavioral strategies (Kunitomo et al., 2013; Satoh et al., 2014). We found that klinotaxis bias was severely impaired in *clh-1(pe)* mutants regardless of cultivation salt concentrations (Figure 6**a**, Figure 6—figure supplement 1**a** and **c**). Klinokinesis bias of the *clh-1(pe)* animals after cultivation at 100 mM NaCl was comparable to that of wild type (Figure 6**b**, Figure 6—figure supplement 1**d**). These results suggest that chemotaxis of *clh-1(pe)* animals to high salt after cultivation at 100 mM NaCl largely depends on klinokinesis. However, after cultivation at 0 mM NaCl, klinokinesis bias was lost in *clh-1(pe)* mutants. Up-regulation of pirouette frequency along with salt concentration increase (dC/dt > 0) was abolished in *clh-1(pe)* mutant animals (Figure 6**b** and Figure 6—figure supplement 1**b**). These results strongly indicate that the defective chemotaxis of *clh-1(pe)* mutants is due to loss of both klinotaxis and klinokinesis biases after cultivation at 0 mM NaCl. The klinokinesis and klinotaxis of *clh-1(tm1243)* mutants were comparable to those of wild type (Figure 6**a**, **b** and Figure 6—figure supplement 1**a**, **b**).

**Figure 6.**
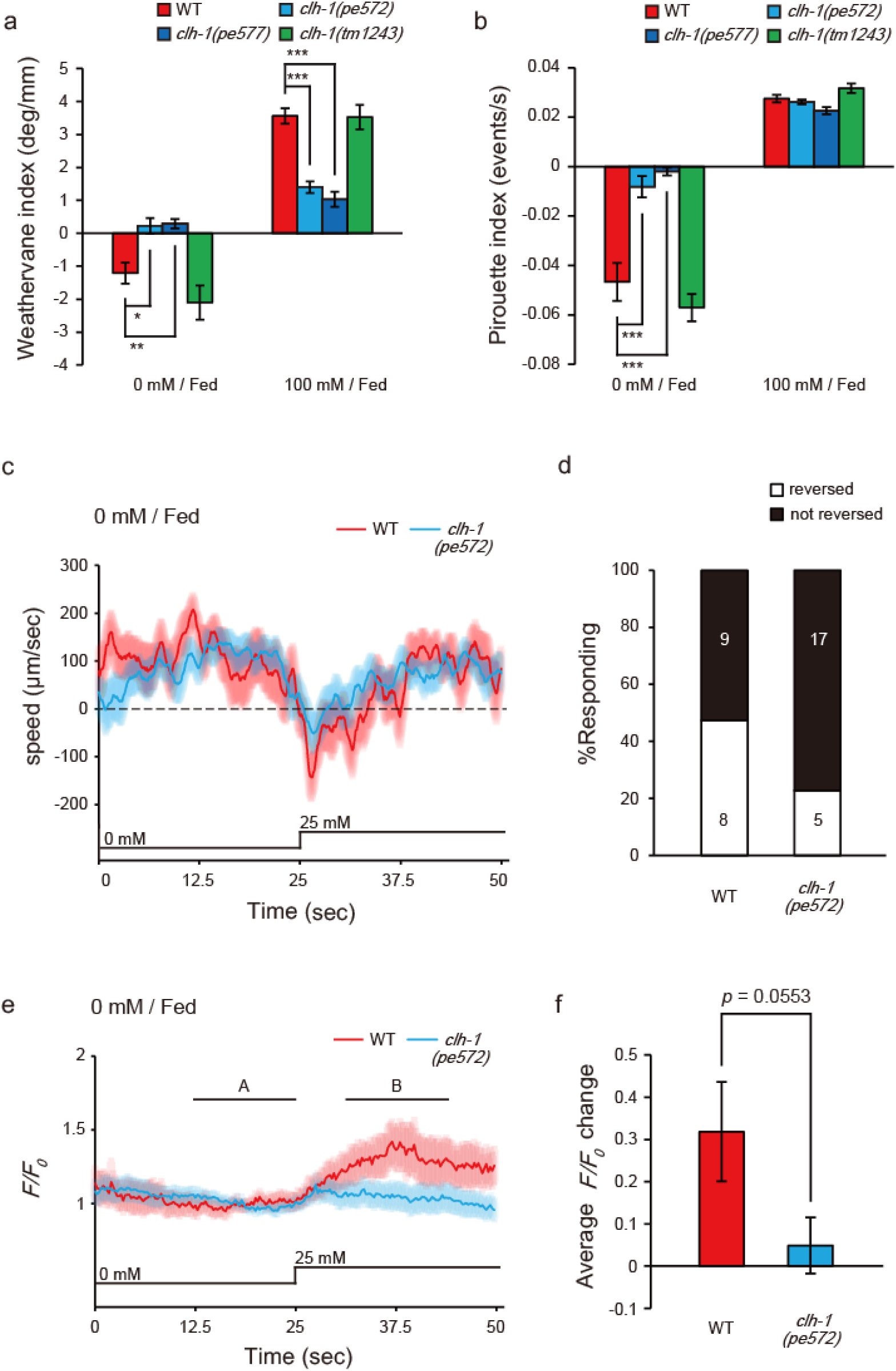
Missense mutations in *clh-1* attenuate both klinotaxis and klinokinesis, also AIB response and reversal in response to salt increase. (**a** and **b**) *clh-1(pe)* mutants show defects in migration bias in salt chemotaxis. Bias of klinotaxis (**a**) and klinokinesis (**b**) are represented by the weathervane index and pirouette index, respectively. In both mechanisms, positive and negative indices indicate migration bias towards higher and lower salt concentrations, respectively. Bar and the error bar represent mean +/− s.e.m., *n* ≧ 18 assays, Dunnett’s test, \*\*\**p* < 0.001, \*\**p* < 0.01, \**p* < 0.05. (**c** and **e**) Averaged locomotion speed of animals (**c**) and GCaMP6s responses of AIB (**e**) after cultivation at 0 mM NaCl in the presence of food. In (**e**), A and B indicate the time points for calculation of Averaged *R/R_0_* changes. NaCl concentration change from 0 mM to 25 mM at 25sec. The shaded region represents s.e.m., *n* ≧ 19 animals. (**d**) Proportion of animals that showed reversal after salt stimulus. Reversal was defined as follows; backward locomotion, whose velocity was less than - 100 µm/sec, was continued for more than 1 second (35 frames). The error bars represent s.e.m., *n* ≧ 19 animals, Fisher’s test. (**f**) Averaged *F/F_0_* change (B - A, see methods). Bar and the error bar represent mean +/− s.e.m., *n* ≧ 19 animals, two-tailed *t*-test with Welch’s correction.

### Reduced salt response of AIB to salt increase in *clh-1(pe572)* mutants

Quantitative analysis of chemotaxis revealed that klinokinesis was disrupted in *clh-1(pe572)* and *clh-1(pe577)* mutants upon increase in salt concentration (Figure 6**b**, Figure 6—figure supplement 1**b**). To further gain insight into the neural mechanism of this phenomenon, we focused on AIB, a postsynaptic interneuron of ASER which promotes sensory stimulus-dependent reversals, the trigger of pirouettes (Piggott et al., 2011; Zou et al., 2018). Because the synapse between ASE and AIB is proposed as the site of plasticity that regulate klinokinesis bias in salt chemotaxis (Kunitomo et al., 2013; Luo et al., 2014; Wang et al., 2017), we hypothesized that the responsivity of AIB may be altered in these mutants. To examine this possibility, we observed salt responses of AIB in freely behaving animals using a microfluidics arena (Albrecht and Bargmann, 2011). Animals were cultivated at 0 mM NaCl with food and stimulated by an up-step NaCl stimulus from 0 mM to 25 mM. Wild-type animals responded to the salt stimulus by slowing down or reversal (the moving velocity becomes less than zero, Figure 6**c** and Figure 6—figure supplement 2). *clh-1(pe572)* mutants also showed reduction in speed, but the proportion of animals that exhibited reversal was relatively smaller than wild type (see legend for detailed criterion of reversal, Figure 6**d**). These results agreed with the klinokinesis defect observed on chemotaxis plates (Figure 6**b** and Figure 6—figure supplement 1**b**). Importantly, responses of AIB to salt up-step correlated well with behaviors. AIB was largely activated upon salt stimulus in wild type, whereas less activated in *clh-1(pe572)* (Figure 6**e**, **f**). Collectively, our results indicate that salt signal mediated by AIB is diminished in *clh-1(pe572)* mutants, and resulted in reduction of turning frequency upon salt up-step after cultivation at 0 mM NaCl.

## Discussion

Here, using genetic, molecular, neurophysiological and behavioral analyses, we showed that the ClC chloride channel CLH-1 is involved in migration bias on salt gradient after feeding. This is probably conducted by, i) maintaining ASER responsivity to repeated salt stimuli that affect klinotaxis, and ii) participating in salt up-step response of ASER and thereby of AIB after cultivation at low salt that contribute to klinokinesis (Figure 7). Repeated activation of ASER synchronized with head swing generates biased klinotaxis (Satoh et al., 2014). Calcium imaging of ASER showed reduced responsivity to repeated down-steps in *clh-1(pe)* mutants (Figure 5**a-b** and Figure 5—figure supplement 1**a**). This result implies that temporal resolution of ASER is impaired in the mutants. Behavior analysis revealed that klinotaxis was actually disrupted regardless of previous cultivation conditions (Figure 6**a** and Figure 6—figure supplement 1**a**, **c**). Our data is consistent with the idea that dynamic [Ca^2+^]_i_ fluctuation in ASER that reflects environmental NaCl concentration change is required for generation of klinotaxis. In addition, *clh-1(pe)* mutants showed klinokinesis defect upon salt increase after cultivation at 0 mM NaCl (Figure 6**b** and Figure 6—figure supplement 1**b**). In agreement with this, ASER response to salt up-step, that is, the *R/R_0_* decay2 of [Ca^2+^]_i_, were reduced in *clh-1(pe)* mutants (Figure 5**a** and Figure 5—figure supplement 1**b**). Furthermore, the response of AIB and reversal behavior upon salt up-step were reduced in *clh-1(pe572)* animals (Figure 6**d**, **f**). Suppression of AIB activity results in reduction of turning frequency (Gordus et al., 2015; Piggott et al., 2011), which is consistent with the idea that inability of AIB to properly respond salt increase gave rise to reduced migration toward low salt regions.

**Figure 7.**
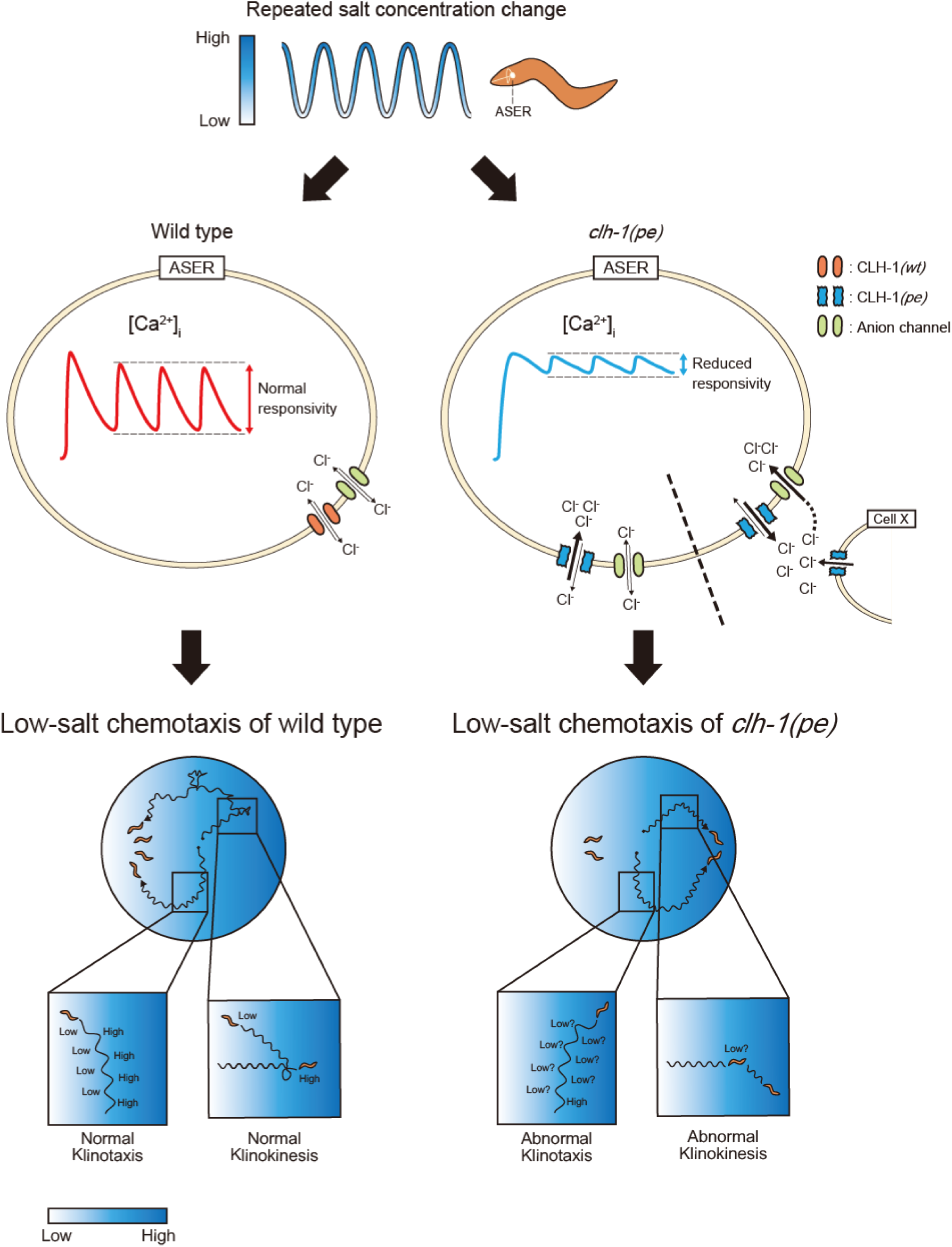
A model for the mechanism of low salt chemotaxis. Animals experience salt concentration changes along with migration on salt gradient, which is depicted by blue waves (top). The salt information is sensed by the ASER neuron and translated into intracellular calcium transients (middle). In the *clh-1(pe)* mutants, calcium responses of ASER to repeated salt stimuli is diminished after cultivation at low salt concentration. This is probably due to increased chloride influx into the cell upon depolarization. Although the mechanism for increased chloride influx is currently unknown, we suggest two possible scenarios; increased extracellular chloride level (right) and increased chloride influx by CLH-1*(pe)* (left). Reduced responsivity of ASER then results in hampered migration bias toward low salt both in klinokinesis and klinotaxis (bottom).

Our results showed that chloride influx into ASER, which was evoked by salt decrease after cultivation at 0 mM, was larger in *clh-1(pe)* mutants compared to wild type (Figure 4**f**, **g**). In general, influx of anion produces outward currents that cause hyperpolarization or prevent depolarization in neurons (Staley et al., 1995). There are several possible mechanisms that could explain increased chloride influx by *clh-1(pe)* mutations (Figure 7, middle). One is a difference in ASER’s transmembrane chloride potential between genotypes. ClC proteins, as well as other chloride transporters such as KCCs and NKCCs, are involved in the excitability of neurons through their homeostatic roles in regulating cellular ionic milieu (Jentsch, 2008; Stauber et al., 2012). Another possible mechanism is elevated anion intrusion via CLH-1*(pe)* channels. ClC-2, the closest mammalian homolog of CLH-1, was shown to be involved in chloride influx into neuronal cells, probably due to incomplete rectification (Ratté and Prescott, 2011; Rinke et al., 2010). In *C. elegans*, CLH-3 inhibit excitability of HSN neurons, which control egg-laying behavior, by directly mediating chloride influx (Branicky et al., 2014). Furthermore, Grant et al. has suggested a possibility that CLH-1 contributes to anion influx with regard to pH maintenance in AmSh (Grant et al., 2015). Since chloride traffic across membranes can affect electrochemical potential of other ions, the altered [Cl^-^]_i_ dynamics might affect cellular [Ca^2+^]_i_ dynamics and excitability. ClC-1, another close mammalian homolog of CLH-1 (37% identity), is involved in excitability of muscle fibers by controlling chloride transportation. Mutations of ClC-1, which are generally attributed to reduction of channel functions, have been identified to cause incomplete repolarization after action potentials, resulting in prolonged electrical activity and muscle stiffness (Adrian and Bryant, 1974; Fahlke et al., 1995; Pusch et al., 1995; Rhodes et al., 1999; Stölting et al., 2014b; Wu, 2002). These reports lend support to the idea that increased chloride influx affect [Ca^2+^]_i_ dynamics of ASER and thereby reduce responsivity of the cell to salt stimuli. These scenarios, together with further characterization of channel properties of CLH-1, needs to be demonstrated in the future study.

In the levamisole resistance test, missense (*pe572* and *pe577*) and the deletion (*tm1243*) allele of *clh-1* showed an opposite phenotype (Figure 1—figure supplement 3**c**). In addition, overexpression of *clh-1(wt)* impaired low-salt chemotaxis; the effect was enhanced in the case of *clh-1(pe)* alleles (Figure 1—figure supplement 2**c** and 3**d**). These results suggest that the *clh-1(pe)* mutations are neomorphic or perhaps hypermorphic alleles. ClC proteins act in homo- or heterodimers (Stölting et al., 2014a). It is not predictable whether CLH-1*(pe)*/CLH-1*(wt)* heterodimer shows mutant-type or wild-type activity. The recessiveness of *clh-1(pe)* mutations (Figure 1**c**) can be explained by assuming that the heterodimer shows wild-type activity.

ASER makes synapses to three first layer interneurons, AIA, AIB and AIY. Activation of AIA and AIY are known to promote forward locomotion, while activation of AIB promotes reversals (Gray et al., 2005; Piggott et al., 2011; Zou et al., 2018). Also, salt response of AIB is markedly changed by salt experience. AIB is activated by salt concentration decrease if previously-experienced salt concentrations were high, whereas it is rather deactivated after cultivation at low salt (Kunitomo et al., 2013; Luo et al., 2014). Here we showed that AIB can be activated by salt increase after cultivation at low salt, which was reduced in *clh-1(pe572)* mutant (Figure 6**e**, **f**). Considering that ASER is glutamatergic (Serrano-Saiz et al., 2013), there may be both excitatory and inhibitory transmission between ASER and AIB. Indeed, AIB expresses the AMPA-type glutamate receptor GLR-1, which mediates excitatory glutamatergic inputs from sensory neurons (Chalasani et al., 2007; Zou et al., 2018). Besides, it has recently been suggested that the glutamate-gated chloride channel GLC-3 and AVR-14 may mediate inhibitory inputs to AIB (Kuramochi and Doi, 2019; Summers et al., 2015). These observations raise the possibility that glutamate response of AIB depends on the electrochemical gradient of Cl^-^ across the membrane as well as the balance of excitatory and inhibitory receptors. In other words, under the condition in which inhibitory glutamate receptors were dominant, AIB could be disinhibited by reduction of presynaptic glutamate release. Growing evidence highlights the importance of chloride homeostasis in the function of the nervous system. In mammalian hippocampal neurons, intracellular chloride is elevated during embryonic development and thereby renders GABAergic transmission excitatory, which is necessary for maturation of synaptic network (Pfeffer et al., 2009). Furthermore, chloride transportation through NKCC1 regulates synaptic plasticity and memory formation in adult hippocampal neurons (Deidda et al., 2015). It will be of interest in the future to determine how the extracellular ionic milieu and glutamate receptors orchestrate the responsiveness of AIB.

In this study we showed that ClC chloride channels function redundantly in salt chemotaxis of *C. elegans*. Food-associated salt chemotaxis was normal in each single (*clh-1(tm1243) II*, *clh-2(ok636) II*, *clh-3(ok763) II*, or *clh-4(ok1162) X*) or double (*clh-2(ok636) clh-1(tm1243) II*) mutants of ClC chloride channel genes (Figure 2**a**, **b**). However, the triple (*clh-3(ok763) clh-2(ok636) clh-1(tm1243) II*) mutation had a marked effect on behavior (Figure 2**b**). Interestingly, all three genes are located very closely on chromosome II (4.08 +/− 0.003 cM for *clh-1*, 3.46 +/− 0.003 cM for *clh-2*, 0.50 +/− 0.000 cM for *clh-3*), suggesting that they are derived by duplication and they might share some evolutionarily conserved functions. The CLH-5 putative anion transporter is also located close to the aforementioned ClC channels on chromosome II: 1.01 +/− 0.007 cM. In addition, mutation of *clh-6(tm617) V*, which encodes a putative anion transporter and whose single mutation had no effect on salt chemotaxis, also gave rise to salt chemotaxis defect in combination with *clh-2(ok636) clh-1(tm1243) II* mutations. Only a few studies so far have addressed functional redundancy of ClC family proteins in an organism (Jeworutzki et al., 2014; Stölting et al., 2014a), and our study provides an insight into functional differences and redundancies of ClC family proteins.

**Figure 1—figure supplement 1.**
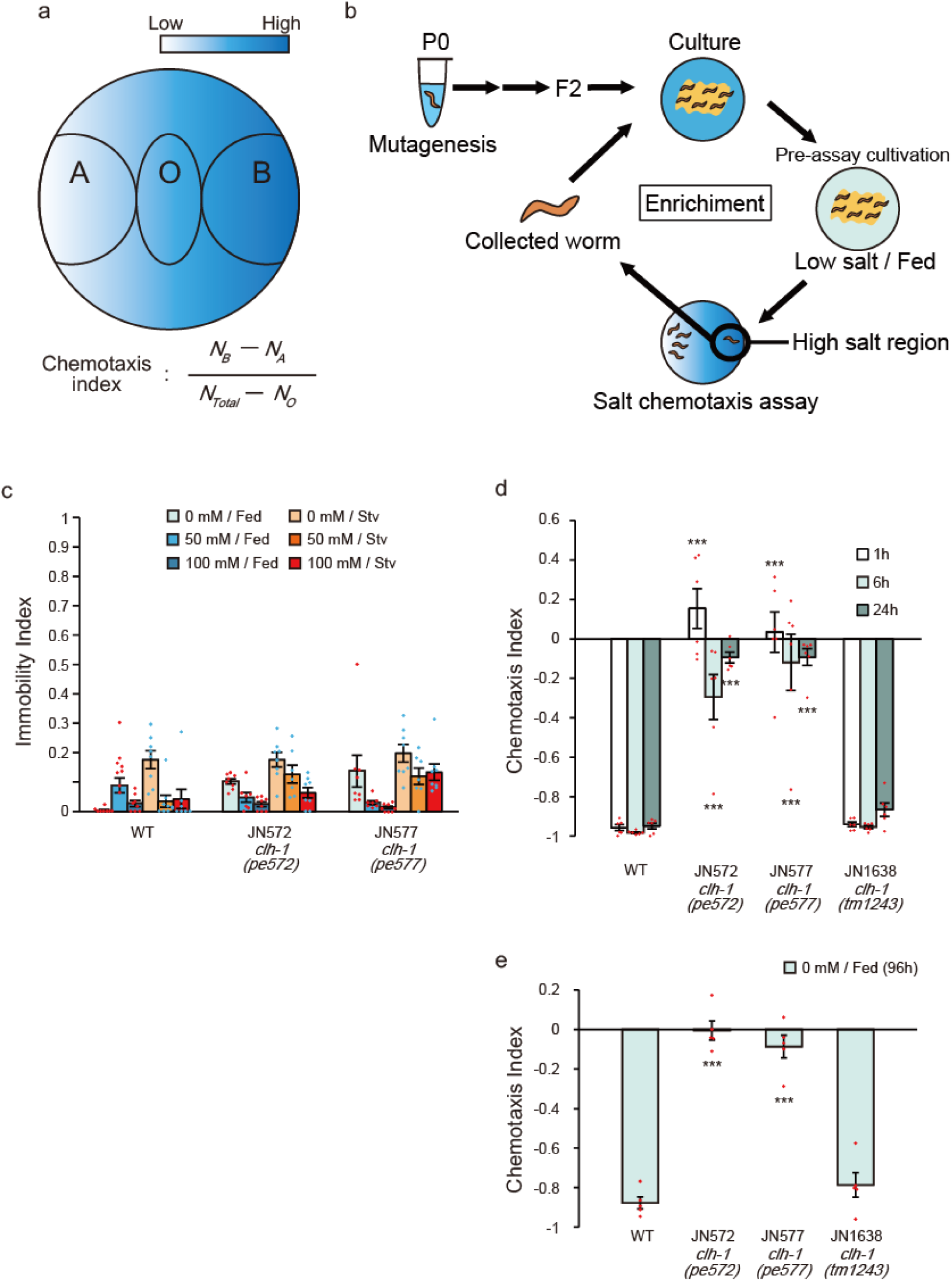
Isolation and characterization of salt chemotaxis mutants JN572 and JN577. (**a**) A schematic diagram of salt chemotaxis assay plate. See methods for details. (**b**) Procedure for forward genetic screening to obtain food-associated salt chemotaxis mutants. Wild-type animals were mutagenized with ethyl methanesulfonate (EMS), and F2 animals were applied for salt chemotaxis assay. Animals that showed defective chemotaxis were isolated and propagated for another round of test to enrich the ratio of mutants. (**c**) Immobility of wild type, JN572 and JN577. See method for details. Dot shows the result of individual trials. Bar and the error bar represent mean +/− s.e.m., *n* = 8 assays. (**d** and **e**) Chemotaxis of wild type and *clh-1* mutants after different duration of pre-assay cultivation. Animals were fed at 0 mM NaCl for 1, 6 or 24 hours (**d**) or 96 hours (from birth until just prior to assay, **e**) and tested for salt chemotaxis assay. Dot shows the result of individual trials. Bar and the error bar represent mean +/− s.e.m., *n* ≧ 6, compared with wild type, Dunnett’s test, \*\*\**p* < 0.001.

**Figure 1—figure supplement 2.**
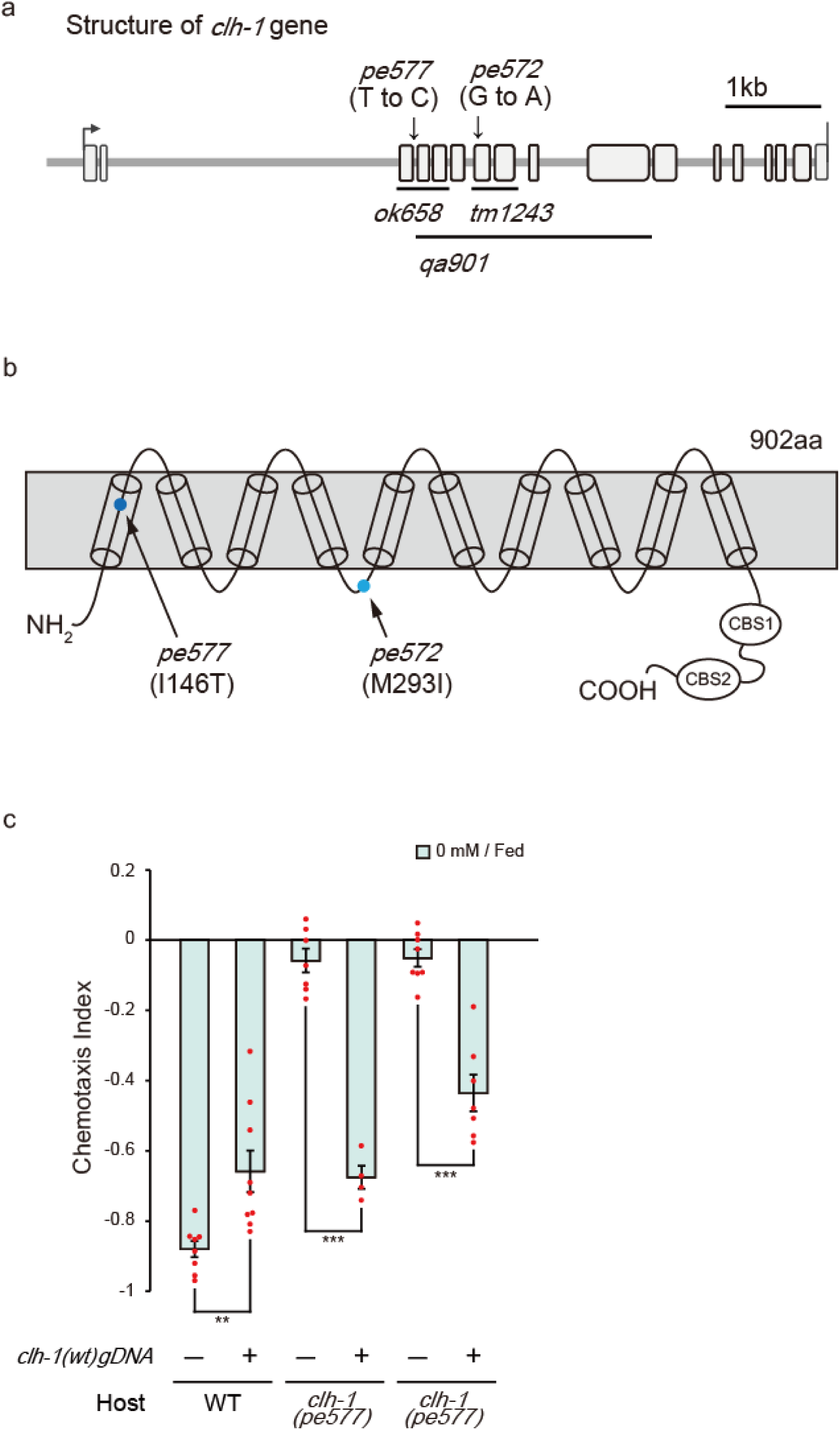
Missense mutations in *clh-1* are responsible for salt chemotaxis defect. (**a**) Gene structure of *clh-1* and the positions of *clh-1* mutations. The positions of *pe572* (T to C) and *pe577* (G to A) are noted by arrows. *ok658* removes exon 3 to 5, *tm1243* removes exon 6 and 7, *qa901* removes exon 4 to 10. (**b**) Predicted protein structure of CLH-1. The positions of *pe572* (I146T) and *pe577* (M293I) are noted by arrows. CBS1 and CBS2 indicate cystathionine beta synthase domains, which are known as protein binding domains. (**c**) Rescue of *clh-1(pe572)* and *clh-1(pe577)* mutants by a *clh-1* genomic DNA fragment. Dot shows the result of individual trials. Bar and the error bar represent mean +/− s.e.m., *n* ≧ 4 assays, Tukey’s test, \*\*\**p* < 0.0001, \*\**p* < 0.01.

**Figure 1—figure supplement 3.**
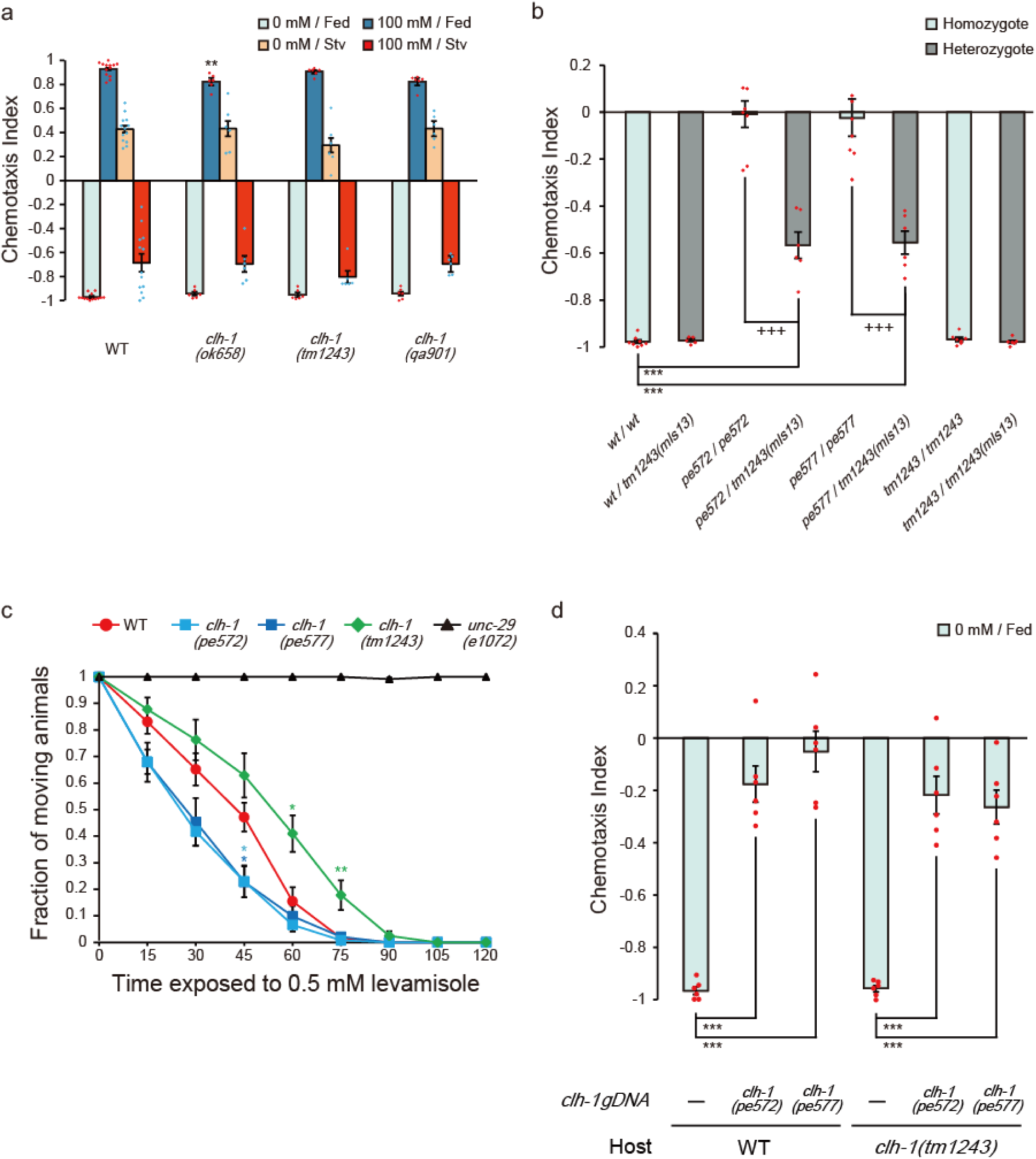
Characterization of *clh-1* mutants. (**a**) Chemotaxis of *clh-1* deletion mutants. Dot shows the result of individual trials. Bar and the error bar represent mean +/− s.e.m., *n* ≧ 6 assays, compared with wild type, Dunnett’s test, \*\**p* < 0.01. (**b**) *pe572* and *pe577* show haploinsufficiency. For generating heterozygotes, hermaphrodites of either wild type, *clh-1(pe572)*, *clh-1(pe577)* or *clh-1(tm1243)* were crossed with *clh-1(tm1243)* males which express GFP in pharyngeal muscle (*mIs13*), and F1 animals were used for the assay. Dot shows the result of individual trials. Bar and the error bar represent mean +/− s.e.m., *n* ≧ 5 assays, Tukey’s test, \*\*\**p* < 0.001, compared with wild-type homozygote. +++*p* < 0.001, compared with each original missense allele homozygote. (**c**) Levamisole resistance test. Graph shows fraction of non-paralyzed (moving) animals at indicated time after placed on 0.5 mM levamisole. Mean +/− s.e.m., *n* = 6 assays, compared with wild type, Dunnett’s test, \*\**p* < 0.01, \**p* < 0.05. Asterisks are colored according to the strain (light blue* for *clh-1(pe572)*, deep blue* for *clh-1(pe577)*, green* for *clh-1(tm1243)* (**d**) Excess genomic DNA fragments of *clh-1(pe572)* or *clh-1(pe577)* recapitulate salt chemotaxis defect in wild type and *clh-1(tm1243)*. Dot shows the result of individual trials. Bar and the error bar represent mean +/− s.e.m., *n* = 6 assays, Tukey’s test, \*\*\**p* < 0.001.

**Figure 2—figure supplement 1.**
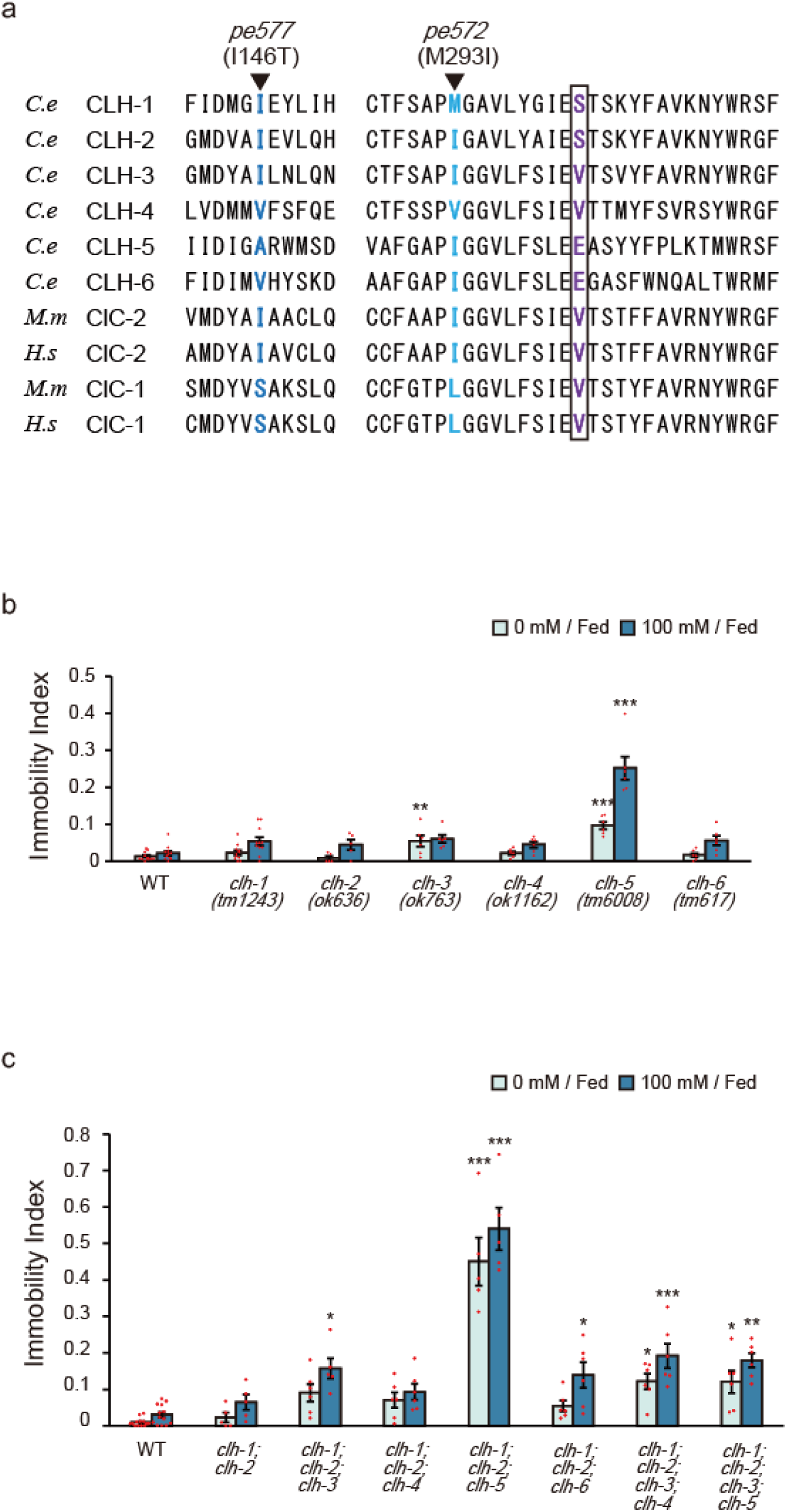
Characteristics of *C. elegans* ClC anion channels and transporters. (**a**) Comparison of *C. elegans* (*C.e*) CLHs, murine (*M.m*) and human (*H.s*) ClC-2 and ClC-1. Amino acids altered in *clh-1(pe)* alleles (deep and light blue) and flanking regions are shown. The “Gating glutamate” residue is boxed, which predicts whether the corresponding ClC protein is an anion channel or a H^+^/Cl^-^ antiporter (Dutzler et al., 2002). Members that carry glutamate at the position is proposed as a H^+^/Cl^-^ antiporter, whereas anion channel if it is valine (V). Although CLH-1 and CLH-2 carry serine (S) residue at this position, they are presumed to be anion channels because they share high similarity with mammalian ClC-2 (Schriever et al., 1999; Nehrke et al., 2000). (**b**) Immobility of *clh* deletion mutants (*clh-1* to *clh-6*). Dot shows the result of individual trials. Bar and the error bar represent mean +/− s.e.m., *n* ≧ 5 assays, compared with wild type, Dunnett’s test, \*\*\**p* < 0.001, \*\**p* < 0.01. (**c**) Immobility of *clh* multiple mutants. Dot shows the result of individual trials. Bar and the error bar represent mean +/− s.e.m., *n* ≧ 5 assays, Tukey’s test, \*\*\**p* < 0.001, \*\**p* < 0.01, \**p* < 0.05, compared with wild type.

**Figure 3—figure supplement 1.**
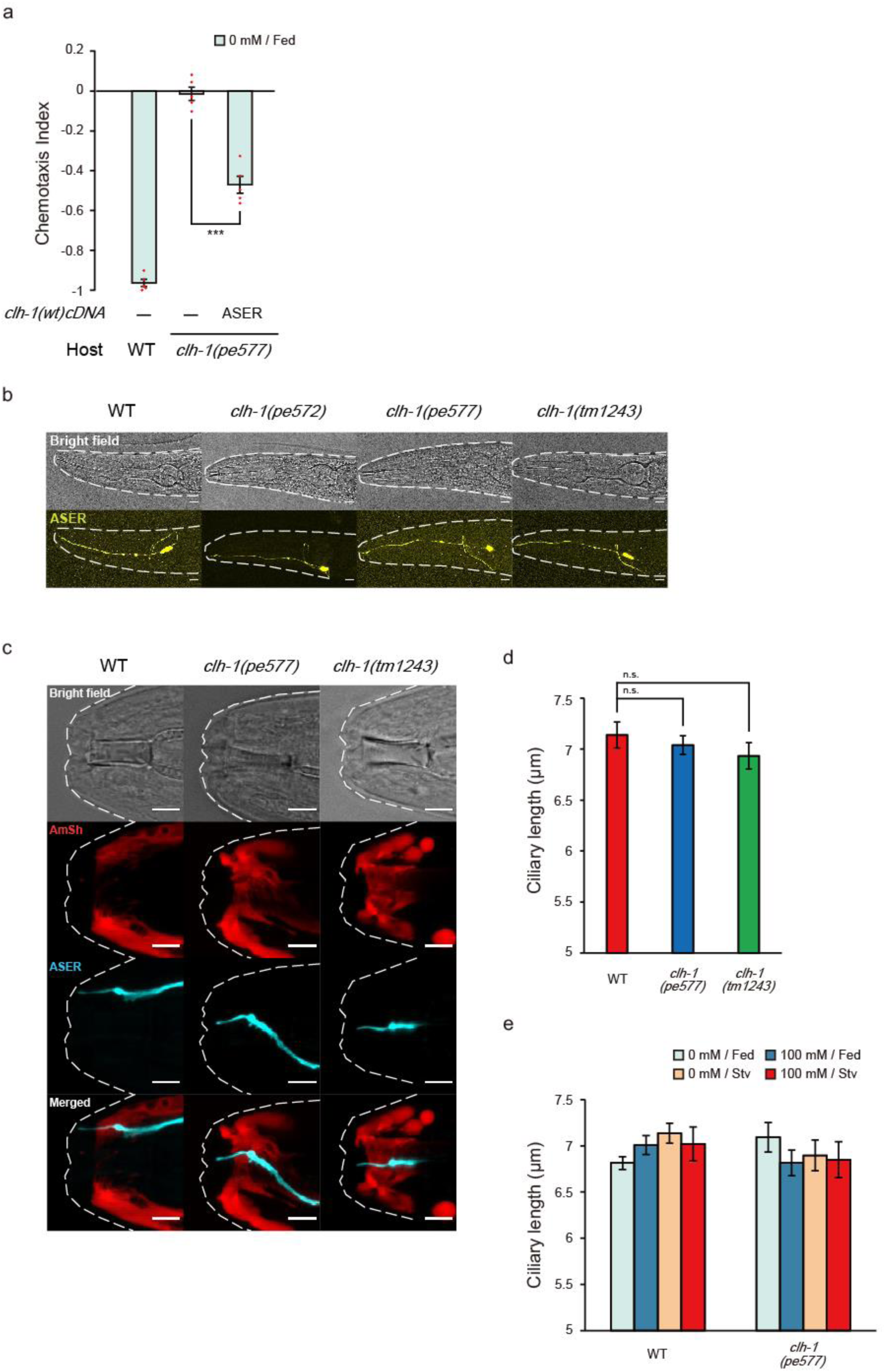
*clh-1* acts in ASER but mutations in *clh-1* do not affect gross morphology of ASER. (**a**) Rescue of *clh-1(pe577)* mutants by ASER-specific expression of *clh-1(wt)* cDNA. The cDNA was expressed in ASER by *gcy-5* promoter. Dot shows the result of individual trials. Bar and the error bar represent mean +/− s.e.m., *n* = 5 assays, Tukey’s test. \*\*\**p* < 0.001. (**b**) Morphology of ASER in wild-type and *clh-1* adult animals visualized by *gcy-5p::GFP*. No gross defect was observed in the mutants. Animals were anesthetized by 10 mM NaN_3_. Bright field (top) and stacked confocal fluorescence images (bottom). Scale bar = 10 µm. (**c**) Confocal images of ASER sensory cilium and amphid sheath glial cells in adult animals. Top; bright field, second top; AmSh (*vap-1p::mCherry*, red), second bottom; ASER *(gcy-5p::mTFP*, cyan), bottom; merged. Animals were anesthetized by 100 mM levamisole. Scale bar = 5 µm. (**d**) Length of ASER sensory cilium in adult animals. Animals were anesthetized by 100 μM levamisole and ciliary length was measured from confocal images. Mean +/− s.e.m., *n* ≧ 16 animals, Dunnett’s test. (**e**) Length of ASER sensory cilium in adult animals after cultivation at the indicated condition. Animals were anesthetized by 100 μM levamisole. Ciliary length was measured from confocal images. Mean +/− s.e.m., *n* ≧ 8 animals, Tukey’s test.

**Figure 5—figure supplement 1.**
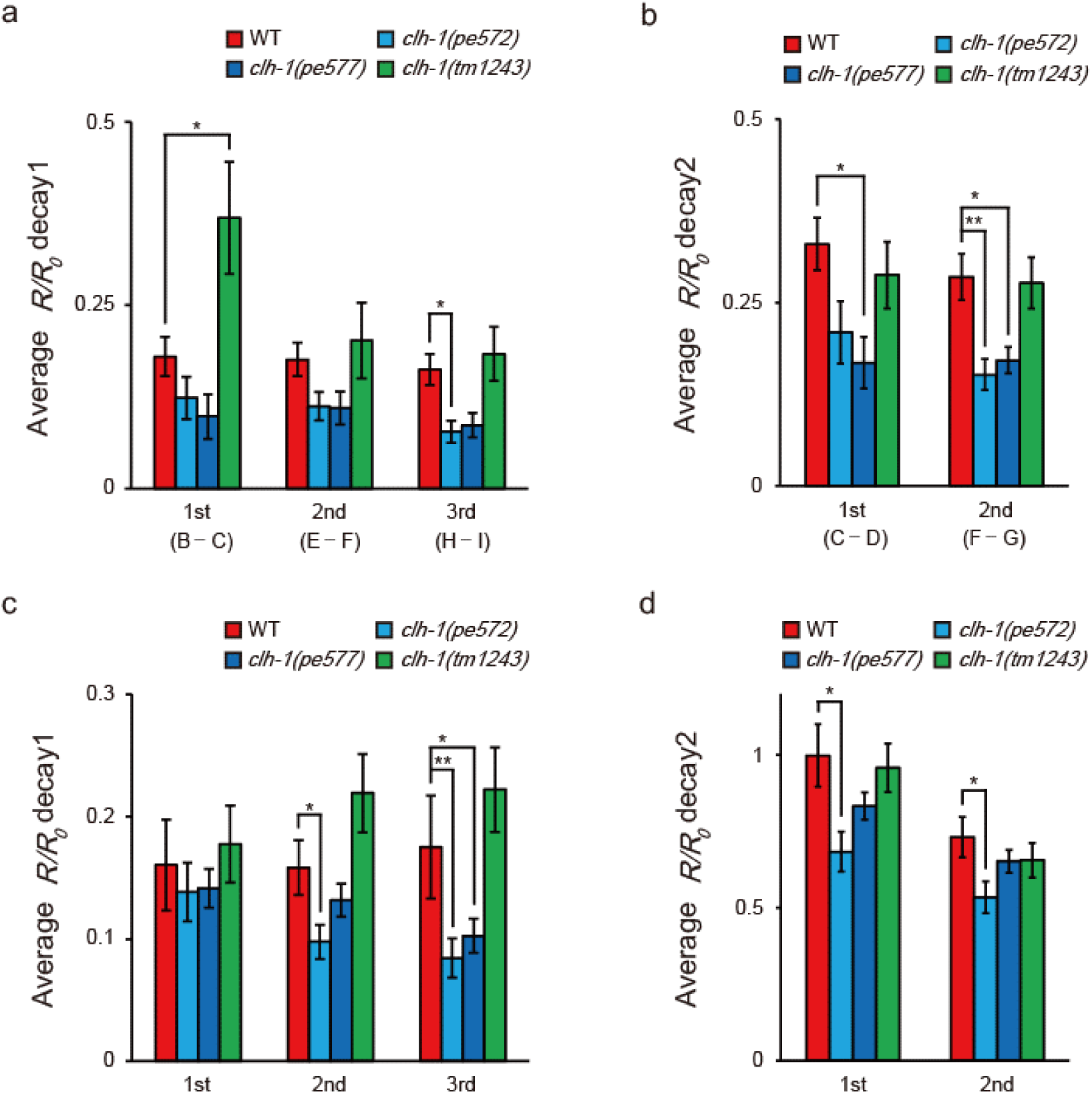
*clh-1* missense mutations affect calcium dynamics of ASER. (**a** and **b**) Decay of ASER calcium response during 25 mM NaCl (**a**) and 50 mM NaCl (**b**). Animals were cultivated at 0 mM NaCl in the presence of food and stimulated by repeated salt concentration change as shown in Figure 5a. Differences of *R/R_0_* intensities at the time points illustrated in figure 5a (B to I) were calculated along with time. Calculation of “decay1” during 25 mM NaCl (**a**) was as follows; B - C for 1st, E - F for 2nd, H - I for 3rd. Calculation of “decay2” during 50 mM NaCl (**b**) was as follows; C - D for 1st, F – G for 2nd. See methods for details. Bar and the error bar represent mean +/− s.e.m., *n* ≧ 16 animals, Dunnett’s test, \*\**p* < 0.01, \**p* < 0.05. (**c** and **d**) Animals were cultivated at 100 mM NaCl in the presence of food. Decays were calculated of as same as described in (**a** and **b**), see methods for details. Bar and the error bar represent mean +/− s.e.m., *n* ≧ 16 animals, Dunnett’s test, \*\**p* < 0.01, \**p* < 0.05.

**Figure 5—figure supplement 2.**
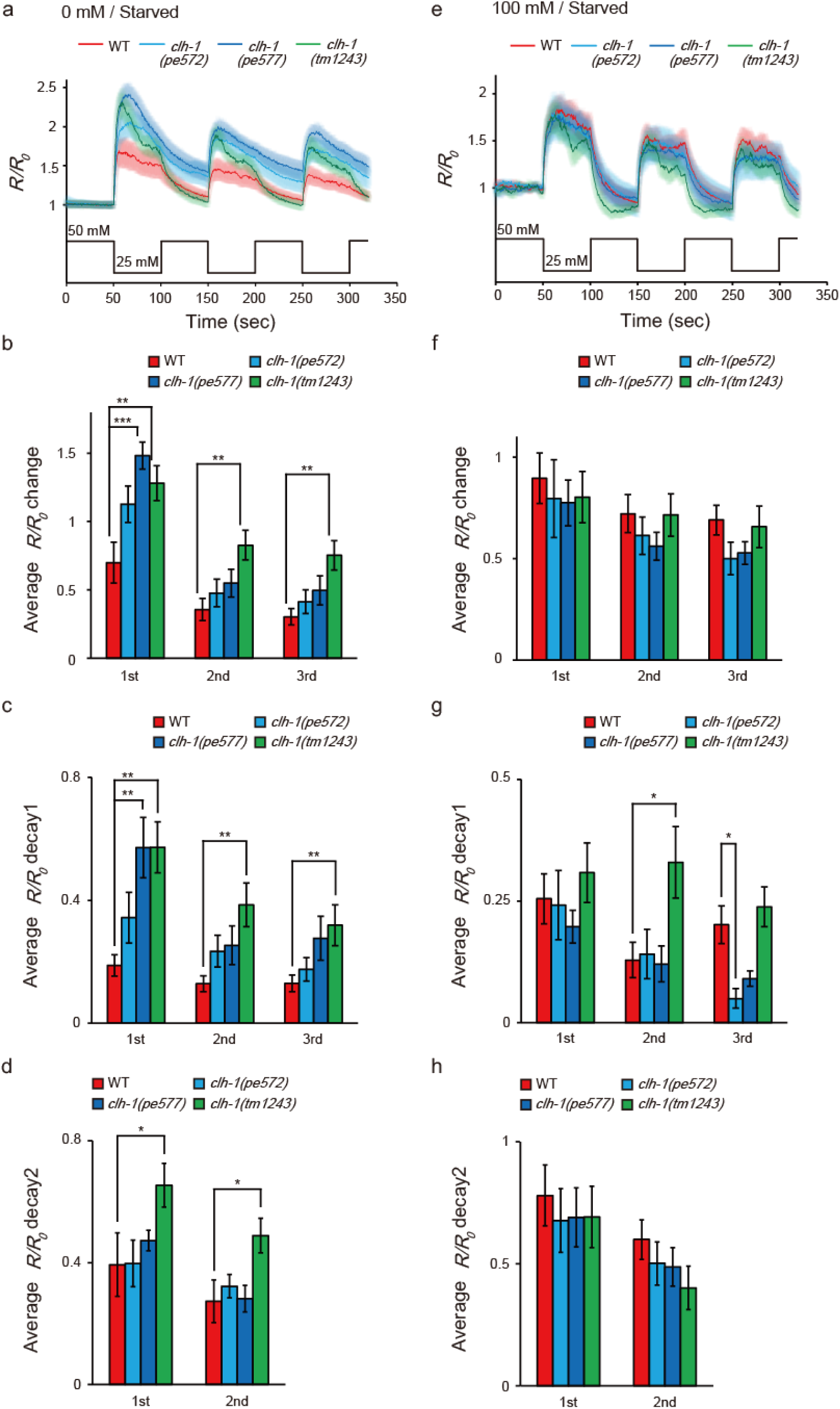
The *clh-1(pe)* mutations less affect calcium dynamics of ASER in starvation-experienced animals. (**a** and **b**) Calcium responses of ASER after cultivation at 0 mM NaCl (**a**) or 100 mM (**b**) in the absence of food and stimulated by NaCl concentration changes between 50 mM and 25 mM. The shaded region represents s.e.m., *n* ≧ 8 animals. (**c** to **h**) Averaged *R/R_0_* intensity changes at each NaCl down-step stimulus (**b**, **f**), decay of ASER calcium response during 25 mM NaCl (**c**, **g**) or 50 mM NaCl (**d**, **h**). 0 mM cultivated (**b**-**d**) or 100 mM cultivated (**f**-**h**). Calculation of *R/R_0_* changes and decays was performed as same as Figure 5a and Figure 5—figure supplement1. See methods for details. Bar and the error bar represent mean +/− s.e.m., *n* ≧ 8 animals, Dunnett’s test, \*\*\**p* < 0.001, \*\**p* < 0.01, \**p* < 0.05.

**Figure 6—figure supplement 1.**
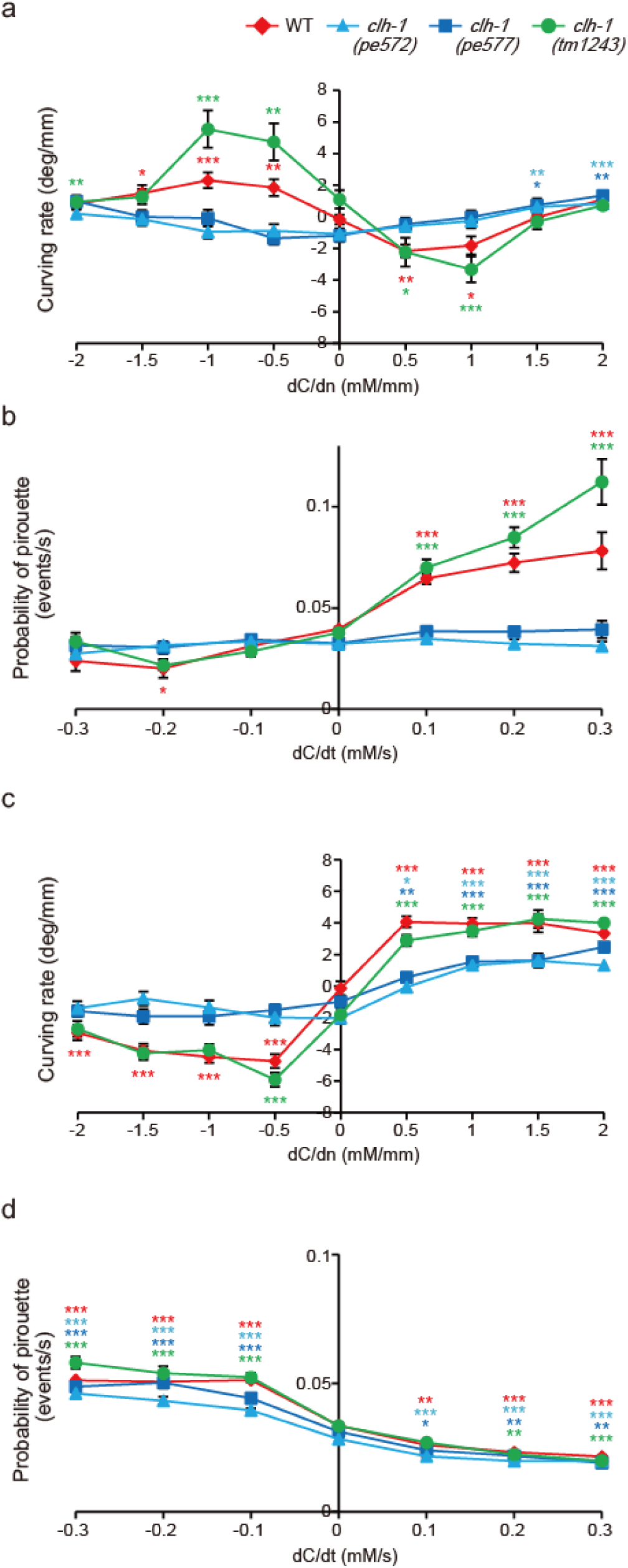
Quantification of navigation behaviors. (**a** and **b**) Chemotaxis after cultivation at 0 mM NaCl in the presence of food. Migration bias of klinotaxis is represented by curving rate (**a**) and the bias of klinokinesis is represented by probability of pirouette (**b**). dC/dn represents spatial gradient of NaCl concentration perpendicular to the direction of centroid movement of the animal. Note that migration biases in both klinotaxis and klinokinesis toward low salt are weak in the *clh-1(pe)* mutants. The error bars represent s.e.m., *n* ≧ 18 animals, compared with the basal weathervane index (dC/dn = 0) or the basal pirouette index (dC/dt = 0), Dunnett’s test, \*\*\**p* < 0.001, \*\**p* < 0.01, \**p* < 0.05. Asterisks are colored according to the strain (red* for wild type, light blue* for *clh-1(pe572)*, deep blue* for *clh-1(pe577)*, green* for *clh-1(tm1243)*). (**c** and **d**) Chemotaxis after cultivation at 100 mM NaCl in the presence of food. Migration bias of klinotaxis (**c**) and klinokinesis (**d**). Klinotaxis, but not klinokinesis toward high salt is weak in the *clh-1(pe)* mutants. The error bars represent s.e.m., *n* ≧ 19 animals, compared with the basal weathervane index (dC/dn = 0) or the basal pirouette index (dC/dt = 0), Dunnett’s test, \*\*\**p* < 0.001, \*\**p* < 0.01, \**p* < 0.05. Asterisks are colored according to the strain.

**Figure 6—figure supplement 2.**
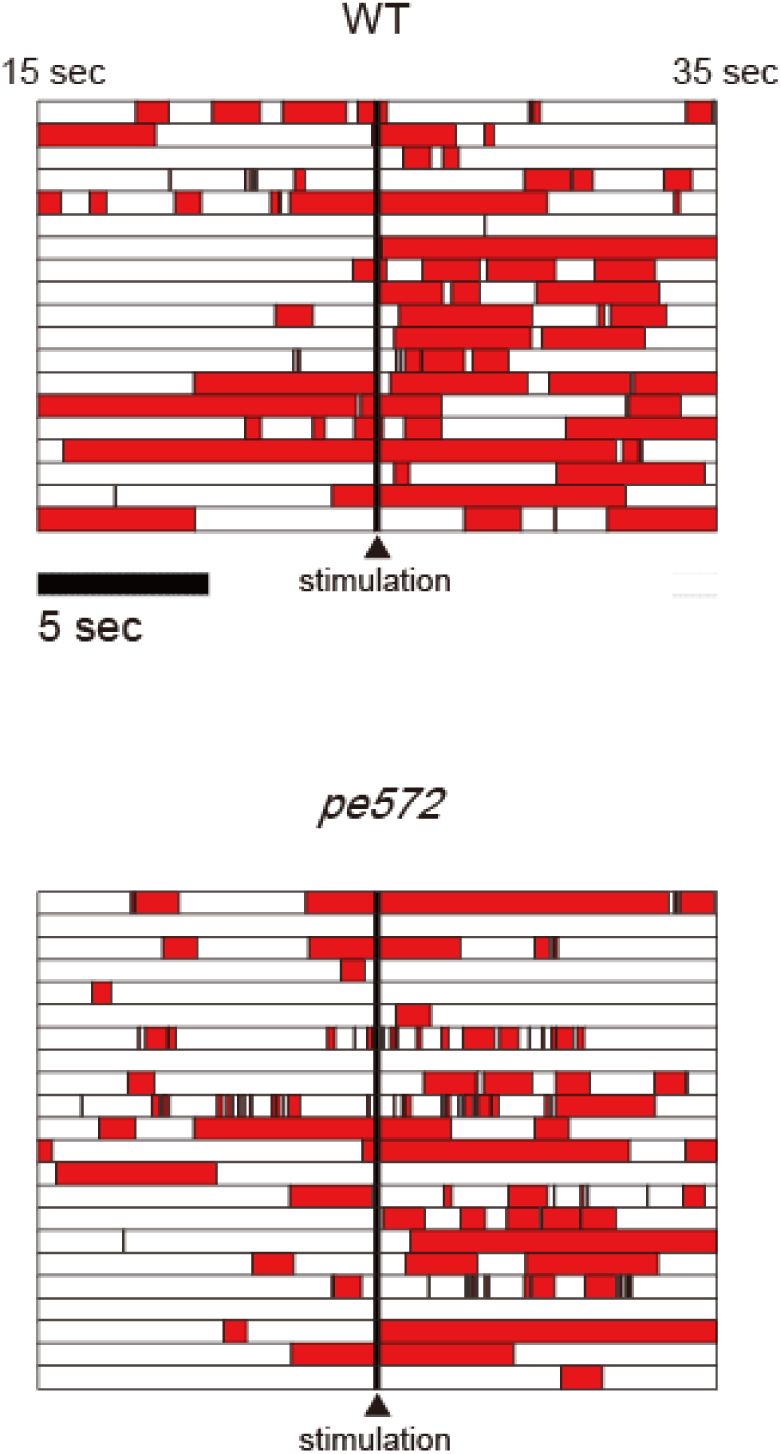
Quantification of reversal behaviors in microfluidics arena. Locomotion of wild type and *clh-1(pe572)*. Locomotion of animal was categorized as either forward (white) or reversal (red). Twenty second recordings, consisting of each 10 seconds of before and after salt up-step stimulation (arrowhead), were aligned. Each row represents each individual. *n* ≧ 19 animals.

## Materials and methods

### *C. elegans* strains and culture

Bristol N2 was used as wild-type *C. elegans*. All mutant strains were outcrossed multiple times with N2. *E. coli* NA22 was used as a food source for behavioral analyses including salt chemotaxis assay. For imaging experiments, OP50 was used as a food source. Strains used in this study are listed in **Supplementary file 2.**

### Behavioral tests

Salt chemotaxis assays were performed as previously reported with minor modifications (Kunitomo et al., 2013). Chemotaxis assay plate was prepared as follows. On top of an agar plate (2 % agar, 25 mM potassium phosphate (pH6.0), 1 mM CaCl_2_, 1 mM MgSO_4_, 8.5 cm in diameter and 1.76 mm in thickness with 10 mL agar), NaCl gradient was created by placing two cylindrical agar blocks (14.5 mm in diameter and 5.3 mm in thickness) that contained 0 mM (position A) or 150 mM (position B) of NaCl in the composition of background plate. Cylindrical agar blocks were removed after 18 to 20 hours, just before assay. Animals were cultivated on regular nematode growth medium (NGM, Brenner, 1974) to young adults, and further cultivated for pre-assay cultivation on NGM plates that contained either 0 mM or 100 mM NaCl in the presence or absence of food for 6 hours. Next, fifty to two hundred worms were collected, washed twice with wash buffer (50 mM NaCl, 25 mM potassium phosphate (pH6.0), 1 mM CaCl_2_, 1 mM MgSO_4_, 0.02% gelatin), then placed at the center of the assay plate. Animals were allowed to run for 45 min. One microliter each of 0.5 M NaN_3_ was spotted to the position A and B so that worms that had reached to these positions were immobilized. The number of worms (*N*) within area A and B (a 2 cm radius from the center of each agar cylinder’s position A and B) and area O (ellipse with radii 20 mm and 10 mm around the start point) as well as the total number of worms on assay plate were counted to calculate the chemotaxis index and immobility index as follows (Figure 1—figure 1 supplement 1**a**).

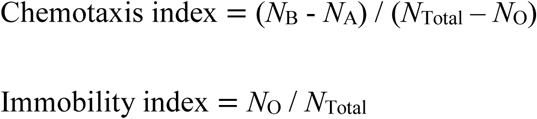

The values of chemotaxis index 1.0 and −1.0 represent complete migration towards higher and lower salt concentrations, respectively, whereas zero value represents no preference for salt concentration (unbiased migration) or a preference for concentration near the central region. Levamisole resistance was determined on agar plates that contained 0.5 mM drug at room temperature. Body paralysis was defined as the lack of body movement in response to prodding by toothpick by visual inspection every 15 min (Gottschalk et al., 2005; Lewis et al., 1980).

### Forward genetic screening and identification of the responsible gene

Wild-type animals were mutagenized with ethyl methanesulfonate (EMS) as described (Brenner, 1974). Progenies were tested for food-associated salt concentration chemotaxis and mutants were selected; worms that showed defective chemotaxis were collected from the assay plate and propagated for testing at the next generation. This process was repeated until F6. We screened approximately 24,000 F2 animals and obtained 7 independent mutants. After being outcrossed with N2, two mutant strains JN572 and JN577 were further analyzed. Other mutants will be described elsewhere.

We mapped the responsible gene for JN572 and JN577, each respectively appeared to be *clh-1(pe572)* and *clh-1(pe577)* (see text), by using single nucleotide polymorphisms (SNPs) between N2 and CB4856 (Fay and Bender, 2006; Wicks et al., 2001) Both of these mutations were mapped between 2.82 (SNP: *WBVar00175127*) and 6.12 (SNP: *WBVar00176673*) on chromosome II. Genome sequences identified a missense mutation in *clh-1* in each strain. We performed rescue experiments using fosmids and genomic PCR fragment of candidate genes. Both mutants were successfully rescued by WRM0612bF09 (fosmid) or genomic PCR fragment of *clh-1*/T27D12.2. Thus we concluded *clh-1* was the causative gene. *pe572* was a G to A transition, whose 5′ and 3′ flanking sequences are TGCACATTCTCGGCGCCTAT and GGAGGTAGGGCTTAACCCTT, respectively. *pe577* was a T to C transition, whose 5′ and 3′ flanking sequences are GATTTTCATCGATATGGGAA and TGAGTATCTGATTCATTGTG, respectively.

### Genotyping

Alleles of each gene locus were verified by PCR using sequence-specific primers for the target sequences. Genotyping primers used in this study are listed in **Supplementary file 3**.

### *clh-1* expression constructs

Full length *clh-1a*/T27D12.2a cDNA clones for wild type, *clh-1(pe572)* and *clh-1(pe577)* mutants were obtained by RT-PCR using sequence-specific primers (5’- GCTAGCCAGGATGGAAGACGCCGTCGTCGT -3’ and 5’- GGTACCTTAGCGGGTTTCGTCATCCG -3’). PCR products were cloned to the pDEST vector (Invitrogen) as a *NheI-KpnI* fragment, and whose sequence was confirmed.

*clh-1* genomic DNA fragments (*clh-1gDNA*s) were prepared by PCR using sequence-specific primers (5’- ATTGCACACATAATTGCGGTAGAC -3’ and 5’- TTGACCCATAAGGTGTAGGCTGC -3’). The nucleotide sequence of open reading frame was confirmed.

### Plasmid construction

Vectors for cell-specific expression in *C. elegans* were generated using the GATEWAY system (Invitrogen). For *clh-1* promoter, a 7.5 kb DNA fragment that contains 4.0 kb upstream of transcription initiation site and 3.5 kb downstream of the first exon of *clh-1a* were amplified from *C*. *elegans* genome (primers: 5’- GCACACATAATTGCGGTAGAC -3’ and 5’- CGCATTTTCTTGAACCCTGG -3’), and cloned into an entry vector, pENTR-1A. For *vap-1* promoter, which specifically expresses in amphid sheath glial cells (Bacaj et al., 2008), we amplified 2.5 kb upstream of the first exon of *vap-1* (primers: 5’- ATTTATAGAAAGTTTCCAAA -3’; 5’- CTGTGAAAATGAACGCACGC -3’). pDEST-*SL2::nls4::mTFP* was generated by ligating the trans-spliced leader sequence SL2, four repeats of the nuclear localization signal, and the fluorescence protein mTFP and cloned into pDEST vector. For pDEST-*clh-1cDNA::mTFP*s, *KpnI-EcoRV* fragment from pDEST*-mTFP* was cloned into the pDEST-*clh-1cDNA* vectors. Superclomeleon (Grimley et al., 2013) was codon-optimized for expression in *C. elegan*s using Codon adapter (Redemann et al., 2011), synthesized, and cloned into pDEST vector. The expression constructs pG-*clh-1p::nls4::mTFP*, pG-*rimb-1p::clh-1(wt)cDNA*, pG-*gcy-5p::clh-1(wt)cDNA*, pG-*gcy-7p::clh-1(wt)cDNA*, pG-*dyf-11p::clh-1(wt)cDNA*, pG-*vap-1p::clh-1(wt)cDNA*, pG-*gcy-5p::Superclomeleon* were created by site-specific recombination between pENTR and pDEST plasmids. Other plasmids have been described previously and details are available upon request.

### Generation of transgenic animals

Expression constructs or genomic PCR fragments were injected at 0.1–50 ng/µl along with a co-injection marker (in many case pG-*myo-3p::venus* or pG-*lin-44p::GFP*, 5-10 ng/µl) and pPD49.26 carrier DNA (a gift from Andrew Fire, up to 100 ng/µl). For comparison among genotypes, the transgene was initially introduced into wild-type background and transferred to other genetic backgrounds by cross. JN2249 was generated by injecting pG-*clh-1p::nls4::mTFP* and *lin-44p::mCherry* into an expression marker strain *Is[rimb-1p::nls4::mCherry; eat-4p::nls4::tagRFP; lin-44p::GFP]*.

### Fluorescence microscopy and measurement of sensory cilium length

Day 1 adults were mounted on 5% agar and anesthetized by 10 mM NaN_3_, or 100 μm levamisole. Images were captured using a Leica HCX PL APO 40×/0.85 CORR CS objective or an HC PL APO 63×/0.40 CS objective on a Leica TCS-SP5 confocal microscope. The length of ASER sensory cilium was measured by simple neurite tracer plugin of ImageJ. All depicted representative fluorescent images were Z-stacked.

### *In vivo* calcium imaging and chloride imaging

Ratiometric fluorescence reporters Yellow Cameleon 2.6 and Superclomeleon were used for calcium and chloride imaging, respectively. We found no obvious defects in salt chemotaxis of the animals that carried these reporters in wild-type background. Imaging experiments were performed as described (Iwata et al., 2011; Kunitomo et al., 2013) with minor modifications. Worms were cultivated on standard NGM seeded with OP50 until adulthood and further incubated for 6 hours at distinct salt concentrations with or without food. Worms were then trapped in a polydimethylsiloxane (PDMS) microfluidics device (Chronis et al., 2007), and NaCl concentration steps were delivered to the nose tip by switching imaging solutions (25 mM potassium phosphate (pH 6.0), 1 mM CaCl_2_, 1 mM MgSO_4_, 0.02% gelatin, NaCl at the indicated concentrations and glycerol to adjust their osmolarity to 350 mOsm). Time-lapse imaging was conducted with a DMI 6000B microscope (Leica) equipped with an HCX PL APO 63x objective (NA, 1.40), L5 filter set (a combination of 430/40nm band-path excitation filter and a 40% transmittance ND filter at 535/40nm dichromatic mirror, Leica), DualView (Filter sets: 505dcxr with 480/40nm and 535/40nm emission filters), and ImagEM EM-CCD camera (Hamamatsu) at two frames per second. All recordings were started 3 min after mounting to stabilize the light response of animals. Fluorescence intensity of the soma was measured. The region of interest (ROI) was tracked by Track Objects function of Metamorph software (Molecular Devices). For each frame, fluorescence intensity of the ROI was calculated by subtracting the background intensity (averaged fluorescent intensity adjacent to the ROI) from the average intensity of the ROI. The ratio of YFP/CFP fluorescence was referred to as *R*. *R*_0_ was defined as the average of *R* over 50 frames (25 sec) prior to stimulation, and the fluorescence intensity ratio relative to *R*_0_ (*R*/*R*_0_) were calculated for a series of images. For traces, *R*/*R*_0_ was averaged for all worms at each time point. The average *R*/*R*_0_ change of Superclomeleon was calculated as the difference between the value of last 10 sec during 25 mM salt stimulus and that just prior to salt concentration change (Figure 4**f**; [averaged *R*/*R*_0_ during B] - [averaged *R*/*R*_0_ during A]). The average *R*/*R*_0_ change of YC2.6 (e.g. Figure 5**b**) was calculated as the difference between the highest value of 10 sec moving average of *R*/*R*0 during 25 mM NaCl stimulus (*R*/*R*_0_ peak) and 10 sec average of *R*/*R*_0_ just prior to corresponding NaCl decrease (e.g. [averaged *R*/*R*_0_ peak of B]-[averaged *R*/*R*_0_ during A] for the 1st, [averaged *R*/*R*_0_ peak of E] - [averaged *R*/*R*_0_ during D] for the 2nd, [averaged *R*/*R*_0_ peak of H] - [averaged *R*/*R*_0_ during G] for the 3rd in Figure 5**a**). Note that the time point used for *R*/*R*_0_ peak differs between individuals because rise speed was different between *clh-1* genotypes. *R*/*R*_0_ decay1 (e.g. Figure 5—figure supplement 1**a**) was calculated as the difference between the *R*/*R*_0_ peak and the average of last 10 sec *R*/*R*_0_ during 25 mM NaCl stimulus (therefore just prior to 50 mM NaCl up-step; e.g. [averaged *R*/*R*_0_ during B] - [averaged *R*/*R*_0_ peak of C] for the 1st, [averaged *R*/*R*_0_ during E] - [averaged *R*/*R*_0_ peak of F] for the 2nd, [averaged *R*/*R*_0_ during H] - [averaged *R*/*R*_0_ peak of I] for the 3rd, in Figure 5**a**). *R*/*R*_0_ decay2 (e.g. Figure 5—figure supplement 1**b**) was calculated as the difference between the average of last 10 sec *R*/*R*_0_ during 25 mM NaCl stimulus and the average of last 10 sec *R*/*R*_0_ during the following 50 mM NaCl period (e.g. [averaged *R*/*R*_0_ during C] - [averaged *R*/*R*_0_ during D] for the 1st, [averaged *R*/*R*_0_ during F] - [averaged *R*/*R*_0_ during G] for the 2nd, in Figure 5**a**).

### Electrophysiology

We synthesized capped CLH-1 cRNAs using T7 mMESSAGE mMACHINE kit (Ambion) and purified by lithium chloride precipitation. cRNA was quantified by spectroscopy. For oocyte preparation, we anesthetized female *X. laevis* in cold MS-222 solution, and excised lobes of ovaries from a small incision made in the posterior ventral side. Oocytes were obtained by defolliculation; incubation of ovaries in 0.2% collagenase (Wako) in modified Barth’s solution (MBSH) for 2 to 5 hours. Oocytes were washed and incubated at 16°C in MBSH with 100U/ml penicillin and 0.1mg/ml streptomycin overnight. Thereafter, oocytes were injected with cRNA mix for a final amount of 40-50 ng/oocyte. Oocytes were incubated in MBSH at 16°C for 2 to 3 days before recording. Currents were measured using a two-electrode voltage-clamp amplifier Oocyte Clamp OC-725C (Warner) at room temperature. Electrodes (0.3–1 mΩ) were filled with 3 M KCl, and then oocytes were perfused in saline with the following composition (in mM): 100 NaCl, 2 KCl, 1 CaCl_2_, 2 MgCl_2_, 10 HEPES, pH 7. For data acquisition and analysis, pCLAMP suite of programs (Molecular Devices) were used.

### Quantitative analysis of animals’ behavior

Quantitative behavior analysis was conducted as described previously with modifications (Jang et al., 2019; Kunitomo et al., 2013). Briefly, animals and chemotaxis assay plates were prepared as ordinary salt chemotaxis assay except that NaN_3_ was omitted. To reduce the chance of collision of worms, only 30–50 worms were placed. Images of whole assay plate were acquired for 15 min at one frame per second. Probability of pirouette and curving rate were calculated as previously described (Jang et al., 2019). Pirouette index was defined as the difference of pirouette probability between negative dC/dT rank and positive dC/dT rank. Weathervane index was defined as the slope of the regression line in relationship between NaCl gradient in normal direction and the curving rate. We exploited data in the range of −0.3 to 0.3 for dC/dT, and −2 to 2 for dC/dn because both pirouette probability and curving rate converged towards zero with high variability in the range of highly negative or positive dC/dT or dC/dn range, probably due to small number of data points at the gradient peaks. We calculated data of each plate, then showed average and s.e.m. in figures. Data from 40 to 540 sec were used to calculate the behavioral parameters because trajectories of worms were highly interrupted by collision of worms before 40 sec.

### Calcium imaging of AIB in freely moving animal

Animals expressed GCaMP6s and mCherry in AIB (JN3329, see **Supplementary file 2**). Worms were cultivated on standard NGM until young adulthood and further incubated overnight on NGM plates with 0 mM NaCl. 20∼25 worms were washed out from the plate, and injected into a PDMS microfluidic device (Albrecht and Bargmann, 2011). After few minutes of acclimation to the PDMS environment under continuous flow of imaging solution without NaCl (see ***In vivo* calcium imaging and chloride imaging** for composition), a 25 mM NaCl up-step was delivered to worms. Bright-field images for locomotion analysis were acquired at 33 frames per second and fluorescence images for AIB activity were acquired at 4 frames per second with a BX51 microscope (Olympus) equipped with a halogen light source (U-LH100IR), a motorized stage (HV-STU02-1, HawkVision), an LMPlanF1 5x objective (NA, 0.13), U-25ND25 (Olympus), a combination of 480/40nm band-path excitation filter and a 25% transmittance ND filter at 505/40nm dichromatic mirror (Leica), DualView (Filter sets: 565dcxr with 520/30nm and 630/50nm emission filters), and a CCD camera (GRAS-c3K2M-C, Point Grey Research). Tracking of particular animal was performed using a Linux-based software (Satoh et al., 2014). Average fluorescence intensity of GCaMP6s and mCherry of the soma were determined by correcting the position of region of interest. After subtracting background, fluorescent intensity of GCaMP6s was normalized by that of mCherry. Average fluorescence intensity over 50 frames (12.5 sec) prior to stimulation was set as *F*_0_ and the fluorescence intensity relative to *F*_0_ (*F*/*F*_0_) were calculated for a series of images. For traces, *F*/*F*_0_ was averaged for all worms at each time point. The *F/F_0_* change in response to stimulation, we calculated the difference of averaged *F*/*F*_0_ between 125 to 175 frames for peak and 51 to 100 frames for baseline (i.e. [averaged *F*/*F*_0_ during B] - [averaged *F*/*F*_0_ during A] in Figure 6**e**). Reversal was defined as backward movement whose averaged velocity was less than 0 µm/s (Figure 6—figure supplement 2, colored in red). In Figure 6**d**, animals those showed obvious reversal response (reversed longer than 35 frames (1 sec) within 105 frames (3 sec) after salt up-step stimulus at velocity less than −100 µm/s) were counted as “reversed animals” to exclude short spontaneous reversals that occur independent of salt stimulus (Figure 6—figure supplement 2, short red bouts).

### Data analyses

The sample sizes were experimentally determined, with referring to those previously reported. Repeats of most experiments were performed in three to six separate days. Statistical analyses were performed using Prism v.5 (GraphPad software, San Diego, CA).

---

**Supplementary file 2:**
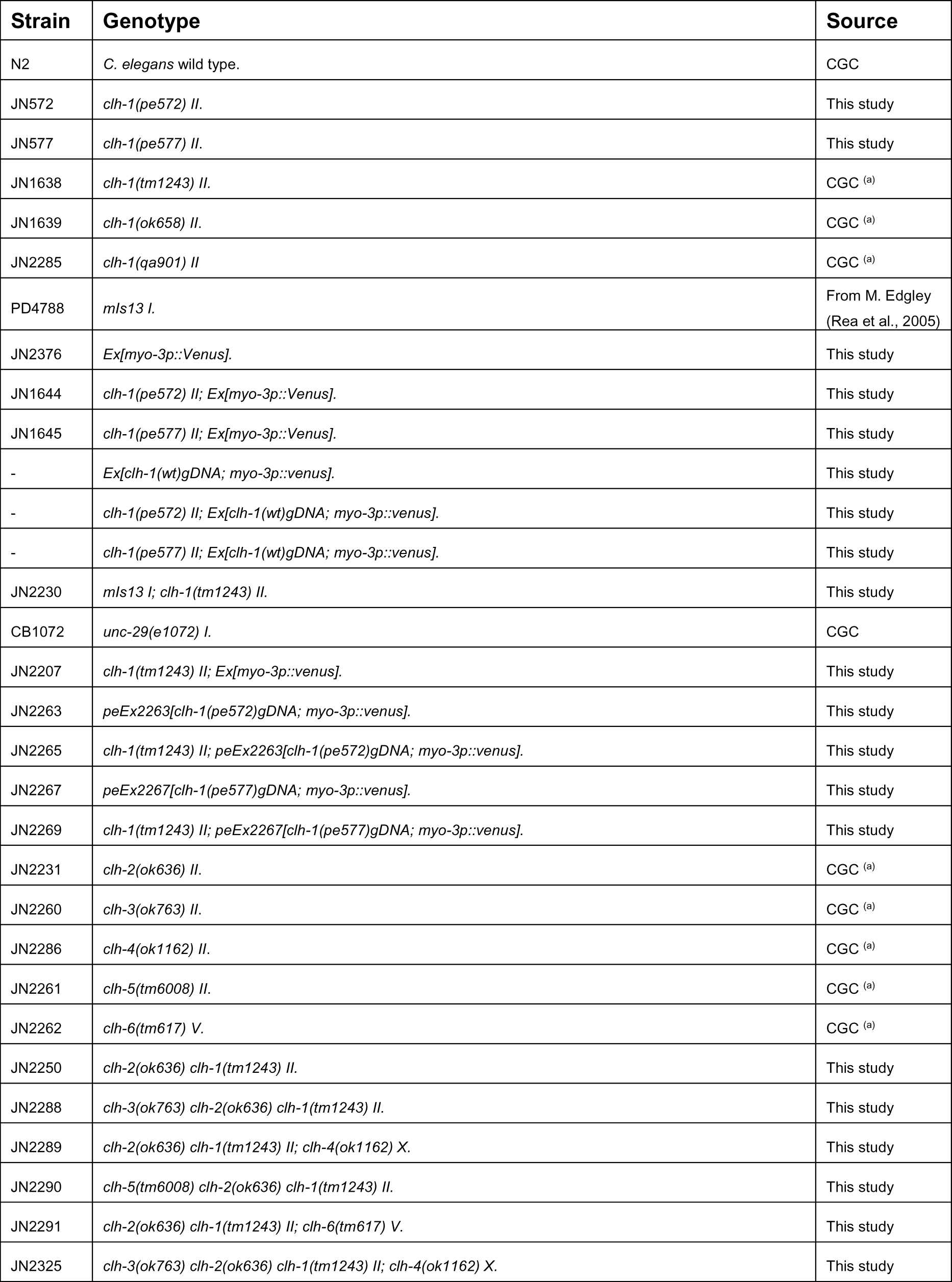
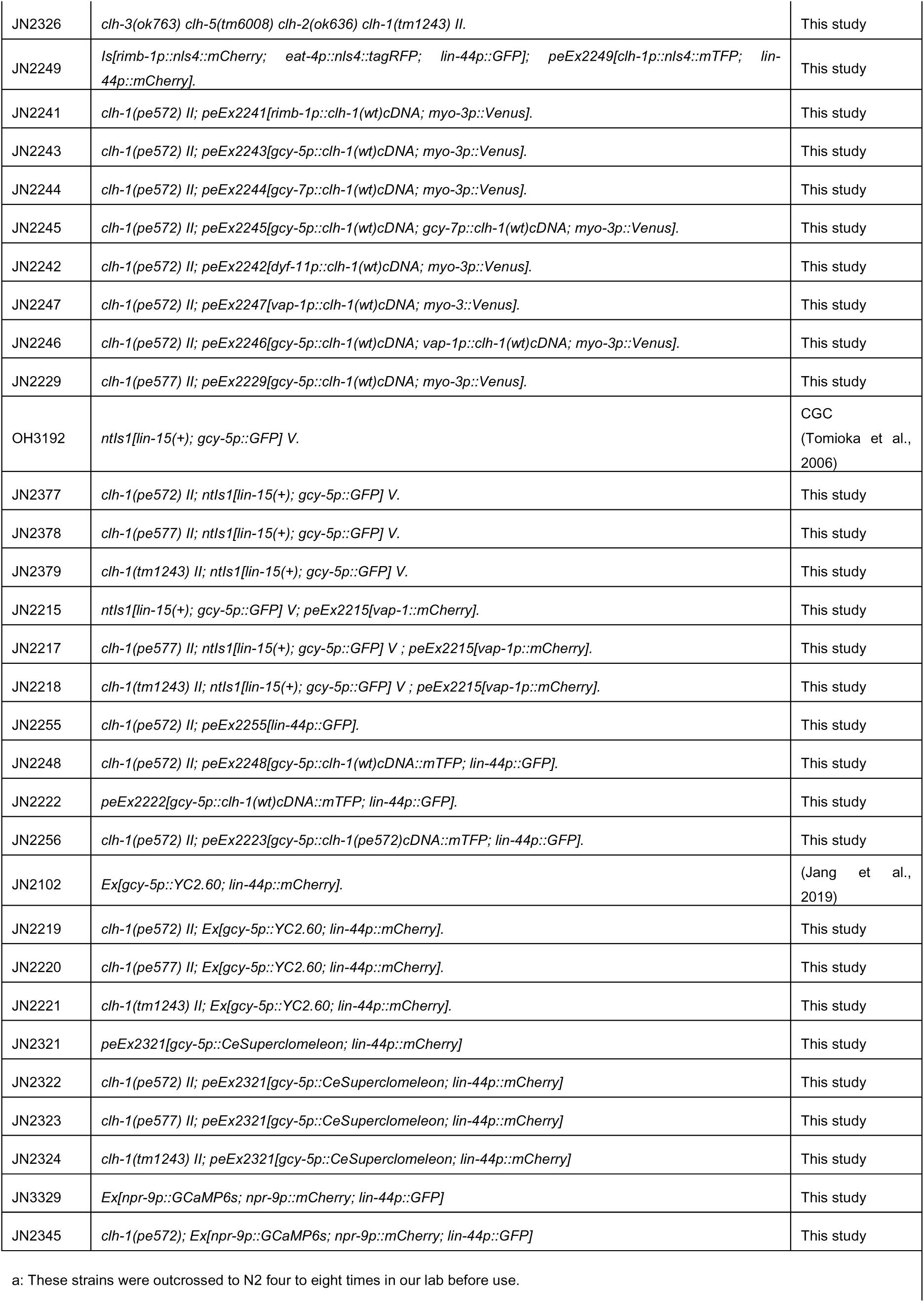
Strains used in this study.

**Supplementary file 3:**
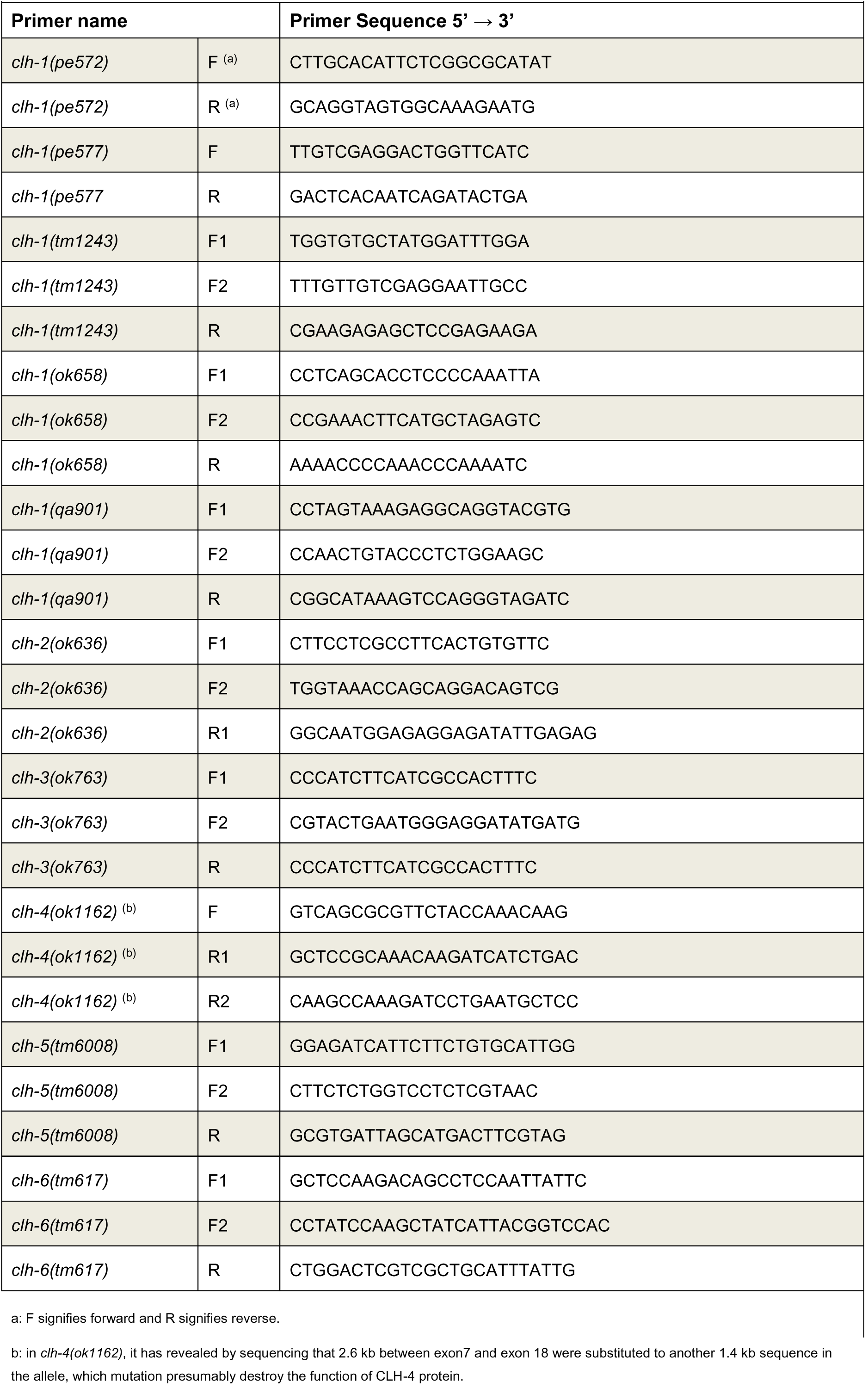
Genotyping primers used in this study.

## Acknowledgments

We thank the Caenorhabditis Genetics Center (CGC) and the National Bioresource Project (NBRP) for strains; Ishihara T. for pDEST-*nls-YC2.60*, Oka Y. and for electrophysiology setup and technical assistance, Chronis N., Albrecht D. and Bargmann C. for design of microfluidics chips. We also thank members of the Iino laboratory members for helpful comments and discussion.

